# Inference of lineage hierarchies, growth and drug response mechanisms in cancer cell populations – without tracking

**DOI:** 10.1101/2025.08.14.669184

**Authors:** Andrea Piras, Federica Galvagno, Letizia Pizzini, Elena Grassi, Andrea Bertotti, Luca Primo, Antonio Celani, Alberto Puliafito

## Abstract

Lineage hierarchies and plasticity regulate development and tissue homeostasis, while diverted lineage dynamics and aberrant phenotypic plasticity are among the causes of incomplete drug response and secondary resistance in cancer. Knowing the dynamics of phenotypically plastic populations is therefore central to understand growth regulation principles and to rationally design therapeutic approaches that might anticipate drug-tolerant states.

Lineage inference however largely relies on single-cell tracking techniques, which are notoriously difficult in complex biological models.

To overcome these limitations, we developed a method to infer active phenotypic transitions in a multi-lineage tumor or clone and to quantify them, solely relying on counting lineage abundances with no pedigree. We demonstrate the effectiveness of our approach to cancer cell plasticity and drug treatment *in silico*. We then perform experiments on cancer cell populations and show that our method correctly predicts growth mechanisms and transition probabilities.

## INTRODUCTION

Tissue development and homeostasis in metazoans rely on the stability of phenotypic identity. In embryonic and tissue development, tightly regulated phenotypic switches are at the basis of the establishment of separate lineages, and hence cellular functions, while plasticity is commonly associated to tissue repair upon insult.

In contrast, cancer cells progressively acquire a cell plasticity which is more pronounced than their normal counterpart, involving rewiring of normal lineage hierarchies or states that are reminiscent of physiological hierarchies^1–8^. Importantly, acquired cancer cell plasticity has also been shown to contribute to therapeutic tolerance and to the emergence of secondary resistance^9,10^. Therefore, it is crucial to relate phenotypic transitions in untreated vs treated tumors in order to reconstruct the effect of the therapy on lineage hierarchies and its impact on drug treatment.

In practice, characterizing phenotypic switches for a set of lineages involves determining all possible transitions, i.e. cell differentiation, self-renewal capabilities, controlled cell death and asymmetric or symmetric cell division. Characterizing a particular lineage behavior *in vivo* requires a number of complex experimental approaches involving clonal analysis and lineage tracing, i.e. observing transitions in real time and/or experimentally tracking the progeny of one (or a few) cells in order to determine the phenotypic traits and lineage of the descendants^11–20^. These approaches however are experimentally challenging even when applied to developmental processes which are biologically well characterized and become often prohibitive when applied to cancer.

Observing lineage transitions in complex biological models is indeed difficult, especially when single-cell level observations are required, due to the necessity of genetically engineering multiple reporters and to follow the dynamics over time. Dynamical measurements are possible by combining the use of fluorescent reporters and that of live microscopy, albeit with a relatively limited throughput^21–23^. Dependence on the progression stage of the tumor, patient variability and unknown and potentially changing genetic or transcriptional markers make these experimental approaches particularly complex. On the molecular side, the advent of single-cell technologies has significantly increased the ability to investigate phenotypic heterogeneity in tissues and several techniques allow us to infer phenotypic trajectories in single-cell datasets (see, for example,^24^) as well as to spatially resolve phenotypic heterogeneity within a tumor. While these techniques can reconstruct a pseudo-temporal order or even estimate probabilities of lineage transitions^25^, they do not have the capability to dissect differentiation mechanisms. For all these reasons, effective quantitative techniques to quantify phenotypic transitional probabilities and that can bypass experimental difficulties are needed.

Classical control theory and continuous model approaches have been applied to infer growth mechanisms in the developing crypt and intestinal tissues^26,27^. Theoretical approaches to decipher the design principles of biological tissue growth have also been followed, combining growth regulation models and lineage dynamics control^28–30^. Specifically, several approaches have been dedicated to lineage reconstruction starting from the observed statistics or dynamics^31–43^. Inference of progenitor types and dynamics can, for example, be performed without witnessing single-cell-level transitions^44^, just by inferring rates from the asymptotic size distribution of clones derived from single cells *in vivo*. Many of these approaches, however, rely on knowing cell histories by tracking, i.e. knowing the complete lineage tree –or pedigree– at all times, or the direct observation of transitions, or alternatively, require to know in advance what transitions are taking place in the experimental model or tissue. Furthermore, additional difficulties are represented by the fact that inferring from small clones or tissue is potentially sensitive to stochasticity and noise, and that transitional probabilities could potentially be changing over time, calling for new approaches based on different theoretical frameworks.

Here we propose a suite of inference methods to obtain single-cell transition probabilities from experimental data of multi-lineage tumors by assuming only knowledge of the number of cells in each lineage, without tracking of sort or the need of knowing the pedigree. Our method bypasses the need of directly witnessing the transitions experimentally and relies solely on the ability of identifying and counting how many cells are found in a particular transcriptional state or ‘lineage’. The proposed approach consists of defining all possible transitions with no prior knowledge and quantitatively inferring the transition probabilities compatible with the experimental observations. This allows to exclude absent transitions when inferred probability is zero.

We present both Bayesian and non-Bayesian, analytical and numerical approaches based on the well-known Bienaymé-Galton-Watson (BGW) stochastic branching processes^45–47^. Such a choice naturally allows to take into account phenotypic switches as well as cell deaths, self-renewal and symmetric and asymmetric divisions and extinction for each clone or cell population. We show that we can infer lineage hierarchies in growing multi-type tumors (such as patient-derived tumoroids) with no prior information. We demonstrate that our approach can unveil the phenotypic mechanism of action of a drug in a multi-type tumor with previously unknown plastic transitions fueling drug-tolerant states. Finally we perform experiments on cancer cell populations and apply our inference methods on experimental data demonstrating the efficacy of our approach.

## RESULTS

### Inference of cell transitions in a BGW branching process

We sought to infer population dynamics in a multi-lineage tumor where in principle many different phenotypic transitions are possible. For each cell these are cell death, inactivity and self-renewal and then all possible phenotypic switches from one to another lineage, symmetric and asymmetric cell division into any other lineage or lineages (fig. 1A). Details on the mathematical model allowing to infer transitions are given in the METHODS section. Given a set of observations 𝒲 representing the experimental data, we consider several scenarios that our approaches can handle (fig. 1B). The case of cell tracking is represented by observations where cell identity and transitions are known at all times, typical of live microscopy. This datasets therefore consists in several repeated series where single cells give rise to different trees that are completely known. This is equivalent to say that the full pedigree is available. A second type of observations is represented by consecutive measurements without cell tracking. Here, we suppose we can count cells of any type at all times without however knowing relations and transitions between them. A third data type is represented by connected end-point measurements, where only the initial and final state are known, as in a fixed end-point snapshot. Last, we consider the case where we have snapshots obtained at different times, which however started from different unrelated ancestors.

**Figure 1.**
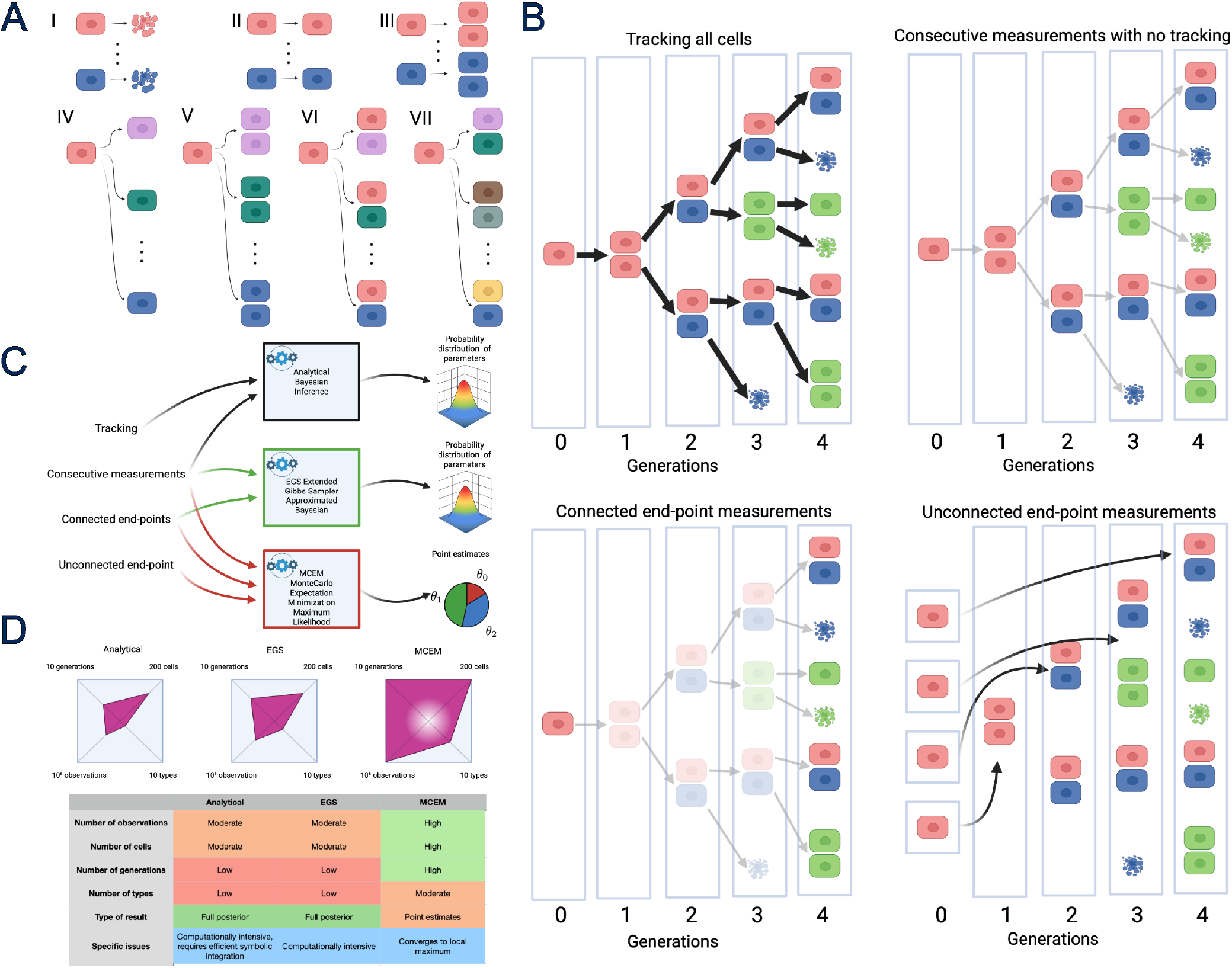
Bienaymé Galton Watson branching processes and inference for cell populations. **A** Illustration of possible transitions in BGW cell population model. **A, I-III** Transitions corresponding to cell death, inactivity and self-renewal respectively for all types. **A, IV** Differentiation into another phenotype. **A, V** Symmetric cell division where the lineage of the offspring is distinct from that of the ancestor. **A, VI-VII** Asymmetric cell division in the two different cases where one of the daughter cells is homotypic with the mother cell (VI) and and where both daughters are different from the mother (VII). **B** Types of data 𝒲 we consider throughout the paper. **C** Types of data input 𝒲 considered throughout the paper and corresponding output for each distinct method developed here. **D** Phase portrait for the models. **Top:** Qualitative phase portrait of each of the methods presented in the manuscript, to highlight the different regions of applicability of those models. The shaded region at the center of the MCEM portrait indicates the fact that MCEM typically requires a moderate amount of data in order to work properly. **Bottom:** Qualitative comparison between the different models and some peculiarity of each of them.

We aim at estimating the probability of all transitions in the cases presented above, particularly when tracking is not available. To this end, we developed a number of different approaches based on the known Bienaymé-Galton-Watson stochastic branching process. In order to handle experimentally relevant situations, we followed three different strategies (fig. 1C). In the first (the analytical method) and second (the extended Gibbs sampler) we employed Bayesian inference, in order to estimate the posterior probability of the phenotypic transitions by knowing the probability of observing a particular observation as a function of the parameters. Details are given in the METHODS section. For datasets with a large number of cells, observations, or cell lineages, Bayesian inference becomes computationally challenging and we resorted to a third computational approach called Monte-Carlo Expectation Maximization. While computationally more efficient, this is a maximum likelihood approach and can only output point estimates for the transition probabilities, rather than entire distributions. A description and comparison of the features of each of these approaches is given in fig. 1C,D.

### Inferring transition probabilities in cancer populations

#### The growth of a two lineages cell population

We begin with the simplest case of unconstrained growth of two distinct non-interacting cell types (a fully developed set of cases for 1 lineage can be found in the SUPPLEMENTARY MATERIAL, sec. 3). A sketch is shown in Fig. 2A. This population structure applies to several different contexts and corresponds to the case where stem cells or progenitors self-renew and give rise to differentiated cells which can only be shed^11,13,26,48^. We assume that the initial population consists of a single cell of type 1 (progenitor) in order to mimic the higher clonogenic potential and the typical situation encountered in cell culture or in clonal development. Note that inference is performed with no prior knowledge, meaning that, for example, differentiated cells are allowed in principle to become stem cells. To tackle the situation, we performed numerical simulations to generate *in silico* data corresponding to fig. 2A that could be fed to our inference algorithm. A set of typical realizations can be seen in fig. 2B.

**Figure 2.**
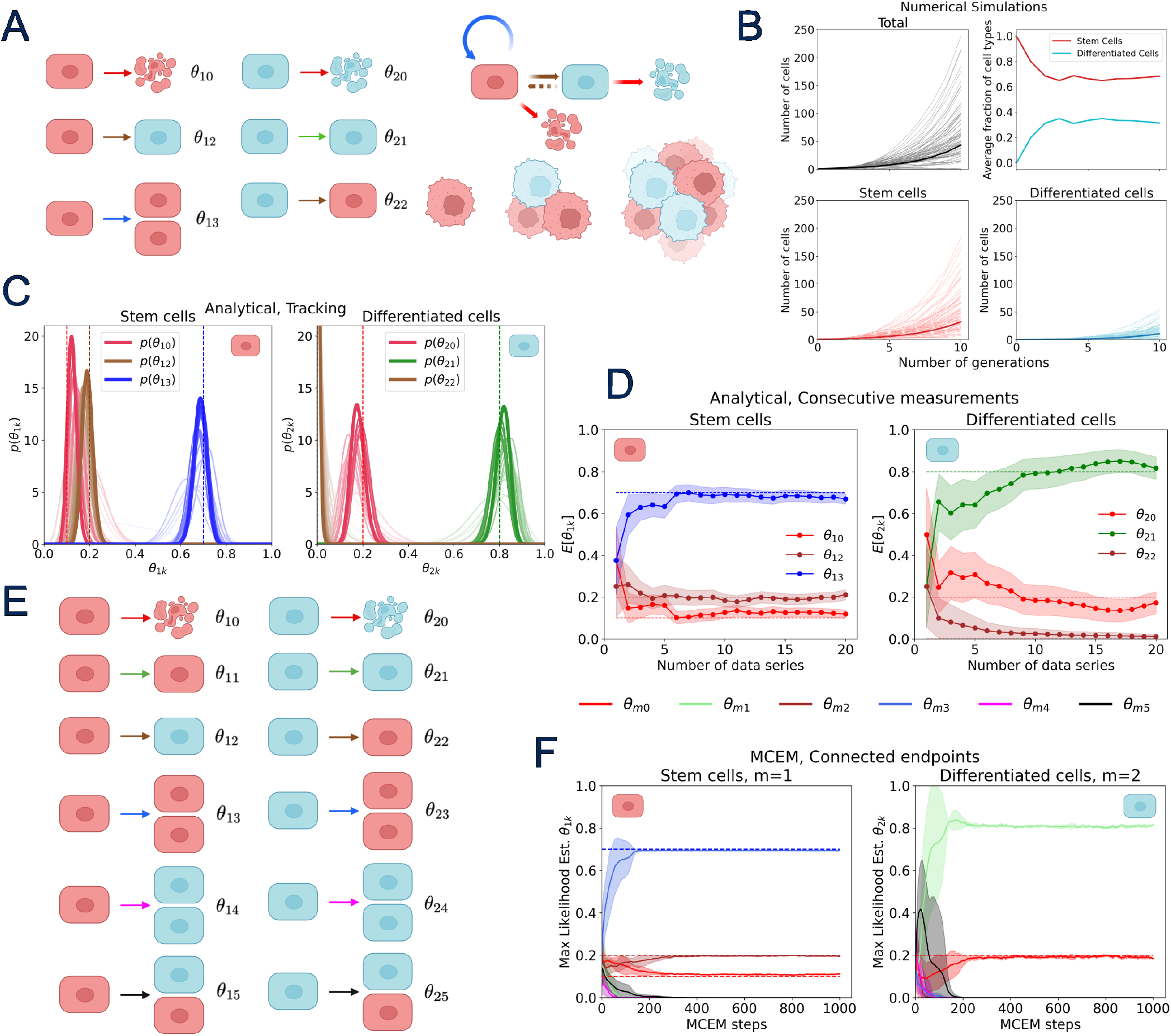
Free growth of 2-types population. **A** (Left) Sketch of the model and parameters used. True values used were 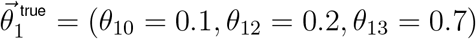 and 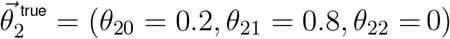. Note that parameter *θ*_22_ is included in inference but set to zero in data. (Right) Sketch of the lineage dynamics of the model, along with a graphical representation of a 2-phenotypes small tumor or tumoroid. **B** Results of numerical simulations of a 2-types BGW branching process with the number of stem cells/progenitors and differentiated cells over the course of generations. The thin light hues refer to single realizations, while thick darker line is the average. The 120 sequences shown were generated starting from 1 cell and lasting 10 generations. **C** Bayesian inference of parameters with cell tracking. In this dataset the likelihoods were 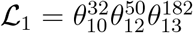 an 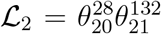. **D** Inference of the parameters by means of the analytical method where tracking information is not available. **E** Full model used with the MCEM method. **F** MCEM estimates with the full model. Endpoint measurements after 6 generations where used as data. Dashed lines represent the true value for each parameter, indicated by a different color, as detailed in the color legend.

#### Results with cell tracking

As shown in the case of 1 cell type (see SUPPLEMENTARY MATERIAL, section 3), the case of tracking reduces to knowing cumulatively all the transitions that occurred in the dataset considered, and this allows to directly write the analytical posterior distribution. Fig. 2C shows the results obtained inferring data obtained with numerical simulations of 1 single progenitor growing for 4 generations and giving rise to structured aggregates. As evident from the plot, the marginal distributions of the inferred values are consistent with the parameter values used to simulated data and converge to the true values when increasing the number of processed data series.

#### Consecutive measurements with no tracking

We then inferred parameters by employing both the analytical method and the EGS assuming no tracking information is available. The feasibility of this approach is demonstrated in fig. 2D, where Bayesian inference is applied recursively. Indeed, our results show that, while data accumulates, the estimates of the parameters converge to the true values. Our analysis also reveals that the level of uncertainty is different in the two parameter sets, due to the relative proportions of the two populations (fewer differentiated cells than progenitors) in each clone and hence in the dataset. It is also worth noting that when prior biological information is unavailable, the full model, which accounts for all possible transitions, must be used. This results in the inference of a relatively large number of parameters, mapping into pronounced variability unless large amount of data is used, which can be a computationally intense task.

Provided that enough data is available, this issue can be addressed using the MCEM algorithm. Notably, this approach remains applicable even when the only available information is the count of the number of cells in each sub-population at the end of the experiment, as typically happens when fixing samples. We demonstrate the suitability of this approach by using 10000 observations representing the size of the two sub-populations after six generations, generated using the previously specified parameter values. This time however, thanks to the better computational performances of the MCEM method, we were able to use the complete model as shown in fig. 2E. The results presented in fig2F show the convergence of the point estimates obtained with the MCEM method to the true values.

We note that our approach can sort out the specific transitions occurring in the dataset by inferring the parameters: death, self-renewal and differentiation of stem cells or progenitors, and death and quiescence of differentiated cells. Therefore such an approach can guide the discovery of lineage hierarchies in a structured population which can then be verified experimentally. It is worth noting that while MCEM is proven to converge, it may only reach a local maximum of the likelihood and not the global maximum. This happens routinely especially in high-dimensional parameter spaces. To effectively reach a global maximum, the algorithm can be restarted multiple times with different initial parameter values. In the absence of the knowledge of the true parameter values, the final likelihood values obtained from different initial parameter configurations can be used to select the parameter set corresponding to the highest likelihood value (see SUPPLEMENTARY MATERIAL, sec. 4).

#### The case of chalones: two lineages with feedback

A natural extension of the two-phenotypes free growth scenario is the case where two interacting phenotypes coexist. Here transition probabilities fuel population dynamics and, in addition, one sub-population can also modulate the growth of the other by feeding back into transition probabilities. Biologically this can occur through paracrine or autocrine signals, such as growth factors or chalones. Similar examples are discussed in ref.^30^ or in ref.^26^, where the balance between asymmetric and symmetric stem cell division is modulated in order to improve developmental times.

To evaluate our ability to infer the feedback mechanisms regulating sub-population dynamics, we selected a specific set of feedback interactions consistent with a scenario where a steady state can be achieved, starting from a single initial stem cell or progenitor. Specifically, we employed the following model for the feedback (fig. 3A,B):

**Figure 3.**
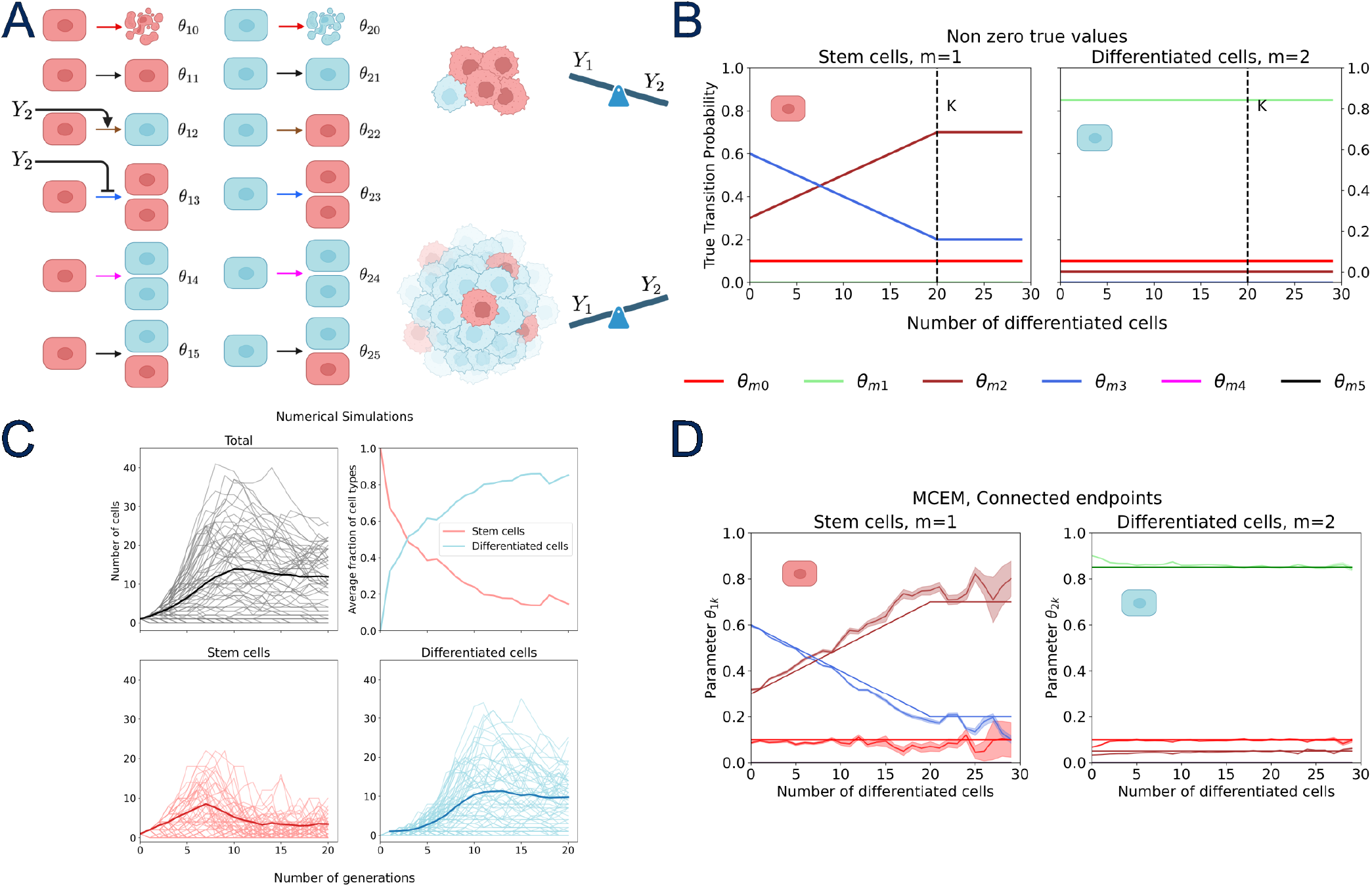
The growth of a tissue with two interacting lineages. **A** Sketch of the model used with the role of feedbacks. **B** Model parameters used to generate data were 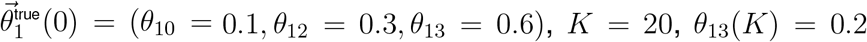 and 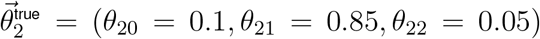. **C** Numerical simulation of the process in panel A: (black) total population size; (red) stem cell/progenitor compartment; (cyan) differentiated compartment. Upper right plot shows the average fraction of stem cells/progenitor (red) to differentiated cells (cyan) over the generations, showing the two distinct regimes of growth. **D** Results of inference from data shown in panels B,C by using MCEM on stem cell/progenitor parameter sets (left) and differentiated parameter sets (right).

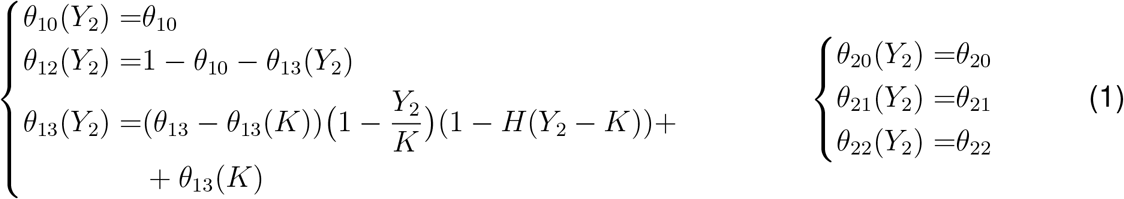

where parameters that are not specified are assumed to be zero, *H*(*Y*_2_ *− K*) is the Heaviside function (*H(x)* = 1 when x *≥* 0 and *H(x)* = 0 elsewhere), and *θ*_13_ *≥ θ*_13_(*K*).

In this model, progenitors (*m* = 1, fig. 3A) can proliferate, differentiate, or die. While the death probability remains constant, both proliferation and differentiation probabilities depend on the population size of differentiated cells. As the latter population grows in each tissue unit, or tumoroid, progenitors increasingly favor differentiation over self-renewal up to certain stable situation reached around a critical differentiated cell population K. Note that differentiated cells can (albeit with a small probability) de-differentiate. A notable consequence is that differentiated cells can replenish the progenitor population in cases where the progenitor compartment has been depleted or eliminated, such as after injury, treatment or targeted ablation^8,49–52^. Data generated with this model exhibit bounded overall growth in both populations, but show two different regimes: one where stem cells prevalently self-renew and stem cell prevail over differentiated cells, and one later regime where a constant ratio between the two types is maintained, in favor of differentiated cells. Results of the simulation are shown in fig. 3C. This situation can be tackled with the MCEM approach. The data containing the number of cells for each phenotype at each generation were used as input for the MCEM algorithm with the aim of inferring the population structure and the relations among the different compartments, i.e. the feedback mechanisms. We note that without any prior knowledge, one should perform inference for each (*Y*_1_, *Y*_2_) pair individually and verify *a posteriori* whether parameters are constant along one direction (in this case *Y*_1_). This approach can be followed using MCEM but requires a large amount of data for each pair in order to obtain reliable estimates. Alternatively, one can hypothesize a dependence only on *Y*_2_ or introduce a prior knowledge suggesting such a dependency. In this case, one can directly group the data of the type (*Y*_1_, *Y*_2_ = *y*_2_) and infer the parameters as a function of *Y*_2_. The comparison between the true and inferred values in fig.3D illustrates this latter approach and demonstrates the capability of our method to uncover the underlying feedback mechanisms. It is also worth noting that inference quality degrades over the number of differentiated cells. Analogously to the previous case, this is due to the number of observations being more and more rare for large *Y*_2_, determining a consequently larger standard deviations of the inferred parameters.

#### Multi-lineage plastic cell populations

As an even more complete example showing the power of our approach we present here the case of a stem/progenitor cell compartment (*m* = 1) which can generate transit-amplifying cells (TA, *m* = 2). These in turn can also expand or transition into differentiated cells (*m* = 3), as shown in fig. 4A-C. The biological reference situation is similar to that of many tissues where however we allow differentiated cells to plastically dedifferentiate^50^. Similar examples are encountered upon injury or ablation in healthy tissues^49,51–54^ or in cancer^7,8^.

**Figure 4.**
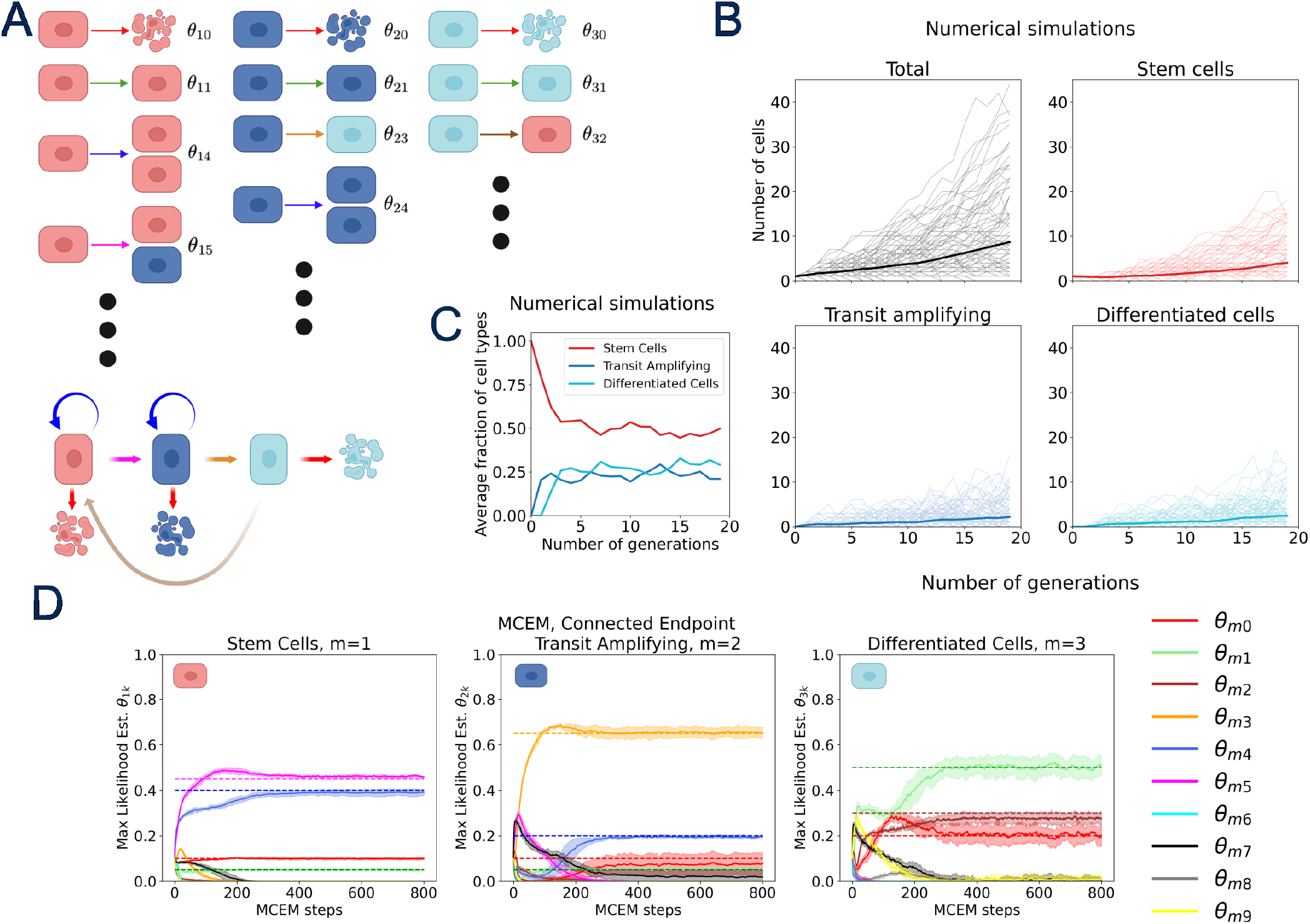
The growth of a multi-lineage plastic tumor. **A** All possible transitions considered are here indicated with their corresponding probabilities. **B** Numerical simulation of the process in panel A: (black) total population size; (red) stem cell compartment; (blue) transit amplifying compartment; (cyan) differentiated compartment. The parameters used for the numerical simulation are 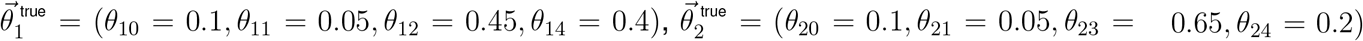 and 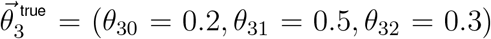, with all other unspecified parameters being zero. **C** Average fraction of stem cells (red), transit amplifying (blue) and differentiated cells (cyan) over the course of each simulation. **D** Maximum likelihood estimates obtained with MCEM on data generated as in panel A for stem (left), TA (middle) and differentiated (right) cells. Colors code for different parameters (transitions) as shown in the legend in panel E.

In order to tackle this scenario, we generated 10000 observations collected after 3 generations starting from one stem cell. On such data we performed inference using MCEM with no prior knowledge and inferred all the 30 parameters of the full model. The comparison between the true values and the estimates obtained exploiting our MCEM algorithm is shown in fig.4D. Once again our approach proves itself successful even in a relatively complex case of a 3 populations hierarchy with no prior knowledge on the transitions.

From an inference perspective, identifying biologically relevant parameters in the full model shown in fig. 1A for 3 distinct lineages, translates into finding 30 parameters in total and therefore requires a substantially larger amount of data with respect to the previous cases. This situation can be attenuated if one introduces a prior knowledge on transitions, for example lack of cell division for differentiated cells, or specifying the type of division (whether symmetric or asymmetric) for stem cells or TA cells.

#### Inferring transitions under drug treatments

In the context of cancer therapy (and in other pathological contexts as well), it is relevant to know how effective is a given drug treatment on cell populations and on lineage dynamics. This problem can be reworded into altered transition probabilities. Indeed, a further, highly relevant, question is what is the phenotypic mechanism of action of a specific drug, i.e. whether it slows down cell duplication, has cytostatic or cytotoxic effects. Within multi-lineage populations this can be impacting on differentiation probabilities or self-renewal probabilities or yet target populations to different extents. In the context of tumor phenotypic plasticity, it becomes relevant to monitor whether a drug treatment is inducing the plastic emergence of new drug-tolerant phenotypes which might fuel relapse or resistance.

However, without monitoring single transitions in a frequentist approach, it is generally difficult to determine the effect of a compound just by looking at cell counts. We therefore address these questions with the tool-set presented above and show that our approach can indeed give answers, even in absence of clear hypotheses or prior knowledge.

#### Treating a one type cell population

We begin by addressing a simple 1-type model, illustrated in fig. 5A,B. This case can be tackled by means of different approaches, presented throughout the paper. First we imagine we can count all occurred transitions, and we therefore write the analytical expression for the posterior. A variation in the observation of the different transitions directly maps into a change of parameters which is captured by analytical inference (as shown in fig. 5C). Results of the inference performed with the MCEM method are shown in fig. 5D, which, as shown previously, can reach larger number of cells and observations. In order to rationalize the effect of the therapy one can exploit all the formalism of the BGW branching processes, calculating extinction probability at a given generation, the mean growth and the ratio between the different transition probabilities. For example, the model shown in fig. 5A is predicted to grow with an average number of cells at the *n*-th generation given by *m* = *Y*_0_(*θ*_1_ + 2*θ*_2_)^*n*^ = *Y*_0_ (1 + *θ*_2_ *− θ*_0_)^*n*^, where *Y*_0_ is the number of cells at the beginning of the treatment. Therefore we can interpret the changes in parameters in the following way: a cytostatic drug will mostly impact on *θ*_1_, while a cytotoxic drug would instead impact the parameter *θ*_0_ by necessarily reducing inactivity and proliferation probabilities and thus affecting the growth rate. For example, a compound leaving the difference *θ*_2_−*θ*_0_ unchanged would not change the overall growth rate but would result in a different standard deviation and would therefore be distinguishable from the untreated case, in contrast with a continuous process. It is worth noting that in presence of parameters inferred in both untreated and treated conditions, it is also possible to evaluate the effect of a drug by measuring the changes in the probability of extinction which can be calculated analytically thanks to the BGW formalism.

**Figure 5.**
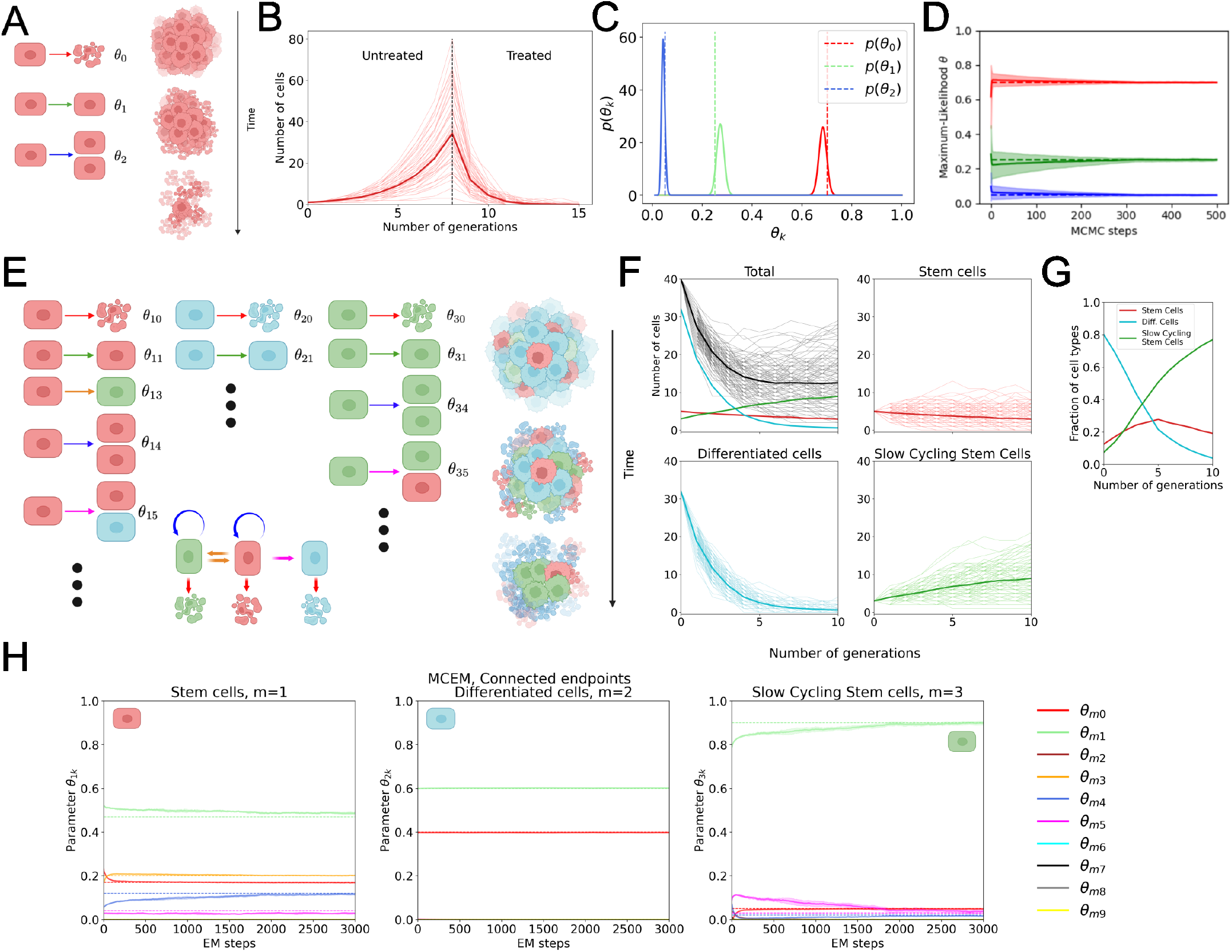
Inferring the effect of a drug on cell populations. **A**,**B** Sketch of experimental settings for a 1-type and 3-types population models for inferring transition probabilities when applying a drug or compound or growth factor. **C** Numerical simulations showing the growth of a cell population with given parameters 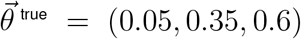 which after 7 gen-erations is treated with a drug with 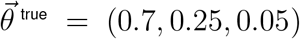. **D** Bayesian inference of parameters in a 1-type setting, corresponding to 12 replicates of data series starting from 50 cells for 3 generations. The likelihood corresponds to 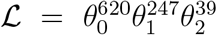. **E** Sketch of the model used to illustrate inference in a multi-lineage treated plastic tumor, made out of 3 phenotypes with stem (red, *m* = 1) and differentiated (cyan, *m* = 2) cells along with a minor population of cells (green, *m* = 3) constituting a population of reserve slow cycling cells that can repopulate the tumor. The parameters used are: 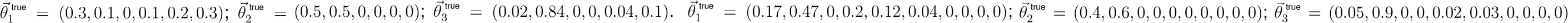. **F** Numerical simula-tions of a treated multi-lineage tumor as in panel E. In these numerical simulations, the tumor is initially composed of 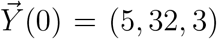 cells. **G** Average fraction of cell types over the course of each simulation. **H** Maximum likelihood estimates obtained with inference performed with MCEM on data generated as in E-G for stem (left), differentiated (middle) and slow cycling stem (right) cells. Colors code for different parameters (transitions) as shown in the legend.

#### Treating a phenotypically plastic multi-type tumor

We then consider the case of a 3-types tumor (fig. 5E-G), where a population of cancer stem cells can self-renew, differentiate or reversibly transition into a slow cycling state^7,55^. Here differentiated cells are post-mitotic and are only progressively shed. We imagine the effect of the drug to be impacting on the death of differentiated cells and cancer stem cells while having only minor effect on the plastic-drug tolerant-slow cycling cells. Under treatment, the overall population shows a significant shrinking, but a small, yet drug-tolerant, population of cells is left behind and is reminiscent of minimal residual disease^56,57^. This situation is clinically relevant, as emerging populations represent a promising target for tumors with incomplete drug response. The effect of the therapy can be seen inferred with the MCEM method in fig. 5H. Here we exploit data collected one generation after the beginning of treatment to infer the impact of the drug on phenotypic transitions. It is worth noting that, as one can expect, such an example is computationally quite demanding as the number of parameters to infer is large. In order to obtain starting conditions for MCEM and ensure convergence, we resorted to the method of moments (or moment matching) (detailed in SUPPLEMENTARY MATERIAL, sec. 8.

Our approach shows that one can determine what phenotypic transitions are particularly involved by the drug treatment under consideration. For example, in the studied scenario, this would unveil a direct impact of the drug on the specific transition transforming slow cycling stem cells into stem cells, which can then repopulate the tumor (fig. 5E). Furthermore, once inferred, the parameters can be leveraged to calculate the extinction probability and this can be used to evaluate the ability to eradicate the tumor in a particular therapeutic setting. All in all, the setting presented here shows the power of our approach in identifying potential drug tolerance mechanisms based on plastically emerging populations and to rationally design population- or transition-targeted pharmacological approaches.

#### Inference from experimental data

We here present the direct application of our inference approach to experimental data for one and two cell types. The need is based rather on assessing the applicability of BGW model to real biological data than on inference itself. BGW models are valid as soon as cell divisions (and other transitions) remain correlated over-time, or occurring within narrow temporal windows. For mammalian cells it was shown that doubling time distributions follow approximately an Erlang distribution with a relatively large shape parameter^58–63^ (a more detailed discussion can be found in SUPPLEMENTARY MATERIAL, sec. 6.) Clonal progenies can therefore be considered correlated for several generations. This is further demonstrated in our experiments.

#### Unconstrained cancer cell growth

In order to directly apply our method to experimental data we seeded the cancer cell line MG-63 at low density in order to observe the offspring of single cells for 96 hours and infer growth parameters. To this aim, cells were transfected with a lentivirus carrying H2B-iRFP670 transgene (fig. 6A). The data onto which we performed inference is a collection of trajectories starting from 1 or 2 cells and followed over 96 hours as shown in fig. 6C. The average doubling time was estimated as *τ*_2_ = (23.1 ± 2.4) hours. First we verified that BGW processes are indeed a good model for the data by looking at the distribution of doubling times within an offspring, as shown in fig. S12A. We found that the relative dispersion is around 10%, confirming that cells remain synchronous for several generations. We applied our inference method to the case when tracking is available, by using the model shown in fig. 6B. To this aim we segmented and counted cells at all frames of our timelapse and detected cell death, identified readily by the absence of movement and by rapid decay of nuclear fluorescence and by evident cell fragmentation. Analogously we could detect cell divisions by looking at binary splitting of cells in the timelapse sequences. By doing so we were able to count the total number of divisions and deaths for each trajectory. Conveniently, when having a number of consecutive tracked observations until the generation *n*_*f*_, the total number of divisions and deaths are related by the relation

**Figure 6.**
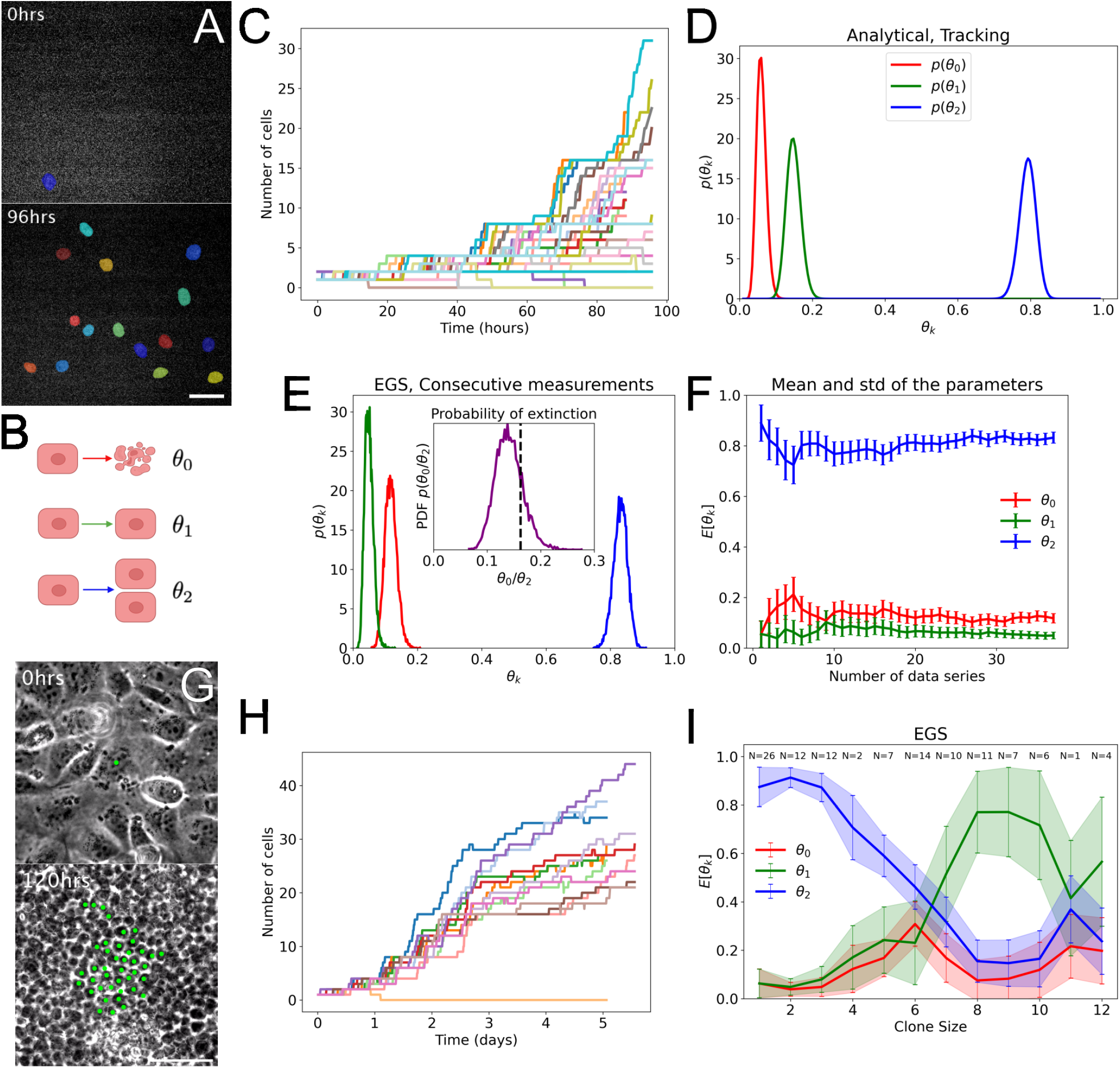
Inferring from experimental data: single type cellular systems. **A** Representative snapshot of MG63-H2BiRFP670 imaged and segmented at two different timepoints cells Scale bar: 50*μm*. **B** BGW model used for inference. **C** Single-cell trajectories monitored for 96 hours. **D** Analytical inference of data shown in panel C with tracking. **E** EGS inference of data shown in panel C. **F** Convergence of mean and standard deviation for inference shown in panel E. **G** Representative snapshots of MDCK cells from ref.^60^ tracked over more than 5 days. Scale bar: 50*μm*. **H** Single-cell trajectories monitored for more than 5 days. **I** EGS inference of data shown in panel H.

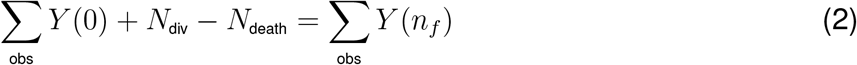

where the sums are intended over all observations. This is because standbys do not change the final number of cells. Therefore, in practice, one can verify the correctness of the death events by directly counting cells and divisions, which with a nuclear marker such as that used in our timelapse is simple.

Further, in order to determine all the transitions observed, one needs to count the standbys. We note that this is also straightforward: it is sufficient to determine the counts at all generations to write:

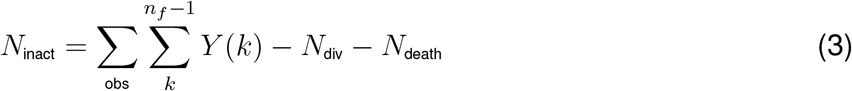

Once the numbers of transitions have been determined one can write the posterior as previously: 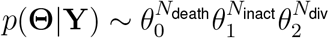. For a set of 37 trajectories lasting 4 generations the results are shown in fig. 6D. It is worth noting that while knowing all transitions for all cells (i.e. tracking) contains all the information to write down the posterior, it is indeed sufficient to count cells at all generations and count all cell division events regardless of the generation in order to write the posterior. When cell divisions are difficult to track and only counts are available we can resort to the other methods developed here. Results obtained with the EGS for the same dataset are shown in fig. 6E,F and are consistent with what obtained analytically, with one difference. Since at this point we used the counting of cells with no tracking, there are many more compatible trees to go from one point to another, resulting in slightly different estimation of the parameters. A more detailed discussion of this aspect can be found in SUPPLEMENTARY MATERIAL, sec. 2.

We note that the inferred parameters can help predicting population properties such as the extinction probability, which can be readily calculated as *p*_*e*_ = *θ*_0_/*θ*_2_ when *θ*_2_ > *θ*_0_ and *p*_*e*_ = 1 otherwise. The probability of extinction at every generation can also be calculated simply by collecting terms in zero-th order in the composite probability generating function of the BGW process (see METHODS, SUPPLEMENTARY MATERIAL and ref.^45^. Notice that in the entire considered database there were 6 clones that went extinct out of 37 which is consistent with the extinction probability calculated theoretically and shown in the inset of fig. 6E.

#### Epithelial contact inhibition

To verify the applicability of our approach to the case of interacting cells we employed data collected in ref.^60^ of MDCK (Madin-Darby Canine Kidney) cells growing onto a 2D substrate (fig. 6G). We tracked the progeny of single cells starting from sub-confluent regimes and followed the offspring for more than 5 consecutive days in order to see the slow-down of cell proliferation (fig. 6H). As before, we estimated the relative dispersion of doubling times within the initial free growth regime and found it to be of the order of 10%, consistently with the previous case of unconstrained growth (fig. S12B). We used these data to infer population parameters. Results obtained with the EGS are shown in fig. 6I. Inferred parameters clearly show that the probability of cell division per generation decreases over time in favor of an increased probability of being inactive with rather constant death probability which is balancing out rare cell division asymptotically. As specified before, no functional dependence of the transition probabilities on density has been assumed here. Results are consistent with what was found in ref.^64^, where cell division is found to be decaying with a power law.

#### Inferring transitions in a multi-lineage tumor

In order to show the ability of our approach to handle multi-lineage tumors we selected a two-phenotypes systems in which, among all theoretically possible transitions, only a few are effectively taking place. To this aim we leveraged the FastFUCCI^65^ system to monitor the cell cycle phase in individual cells (as shown in fig. 7A). We reasoned that by assuming no prior on the possible transitions as usual, we would recover the only possible transitions observed in the cell cycle, i.e. one S/G_2_/M cell to two G_1_ cells (green to two red), and one G_1_ cell to one S/G_2_/M cell (red to green), all the other transition probabilities involving type change being zero (fig. 7B). These have the same phenomenology as a system in which some kind of progenitor can divide by giving rise to two differentiated cells which can revert to the progenitor state.

**Figure 7.**
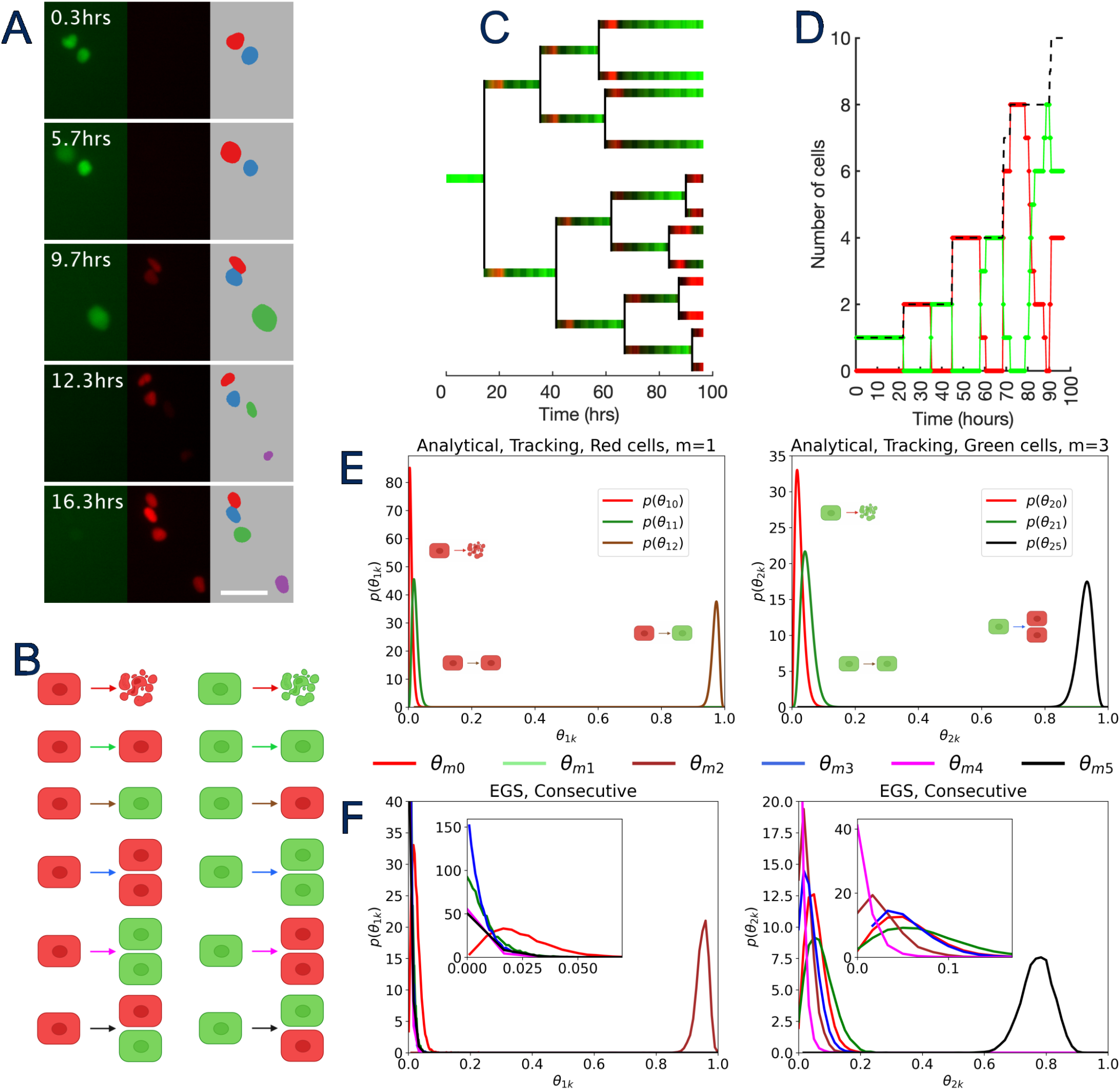
Inferring from experimental data: multi-types cellular systems. **A** Representative snapshots of MG63-FastFUCCI along with cell segmentation. **B** Sketch of the BGW model used for inference. **C** Single-cell trajectory for one cell reconstructed and plotted over the course of the timelapse. Color shades are real fluorescent data for each cell. **D** Data series for a single green (S/G_2_/M) cell evolving into a small clone. Dashed line corresponds to the total number of cells, while red and green count the number of G1 and S/G_2_/M cells respectively. **E** Analytical inference of 16 trajectories with tracking. **F** EGS inference on the same trajectories as in panel E.

First we verified that even in this case data could be modeled with a BGW branching process (fig. S12D). In order to apply our method we built a dataset by segmenting cells and assigning a class (either green or red) by calculating the respective fluorescence intensities (details are given in the METHODS section). A sample of analyzed trajectories is shown in fig. 7C,D. By performing tracking on 20 trajectories we were able to infer parameters, as shown in fig. 7E. As shown by the results, inference performed with all transitions shown in fig. 7B identifies only the expected transitions, i.e. heterotypic cell division where a green cell divides into two red cells and type switch from red to green. In order to illustrate the power of our approach, we performed inference assuming only consecutive measurements, i.e. observing only the counts of each type at each generation without knowledge of the pedigree. Results are shown in fig. 7F and are compatible with the data from tracking. Note that, as previously noted, data from consecutive measurements are less informative than data from tracking (see SUPPLEMENTARY MATERIAL, sec. 2).

## DISCUSSION

In this paper we address the problem of identifying existing phenotypic switches and their probability of occurrence within a population when cell tracking is not necessarily available. Our approach relies on the hypothesis that biological data are reasonably well described by a Bienaymé-Galton-Watson branching process. BGW-based models allow to describe a diverse set of branching processes, they are suitable for different extensions, they allow analytical calculations and can be easily numerically simulated. Such aspects make our approach conceptually very clear and also easily generalizable to other contexts (one above all: epidemiological models^66^). In the context of cancer and cancer therapy we showed how our approach can be used to infer phenotypic transitions in a multi-lineage tumor, and this can of course be applied to normal tissues as well. An interesting application of our method, outlined here, is that of identifying the mechanism of a drug or compound or growth factor on cell populations. It is indeed possible to compare cell populations under treatment and untreated and identify which transitions are mostly affected by the drug.

We have shown that our inference approach is able to quantitatively determine all transition probabilities considered in a general model with no need to track single cells and in certain conditions even with knowledge of endpoint counts only. Such a feature is particularly relevant in the context of cancer phenotypic plasticity where lineage hierarchies might be very different from normal tissue and no strong prior might be available. This applies in particular to the notion of plasticity itself, i.e. reversibility of –normally unidirectional– transitions, which can be determined quantitatively.

The inference technique presented here exploits the statistical distribution of observed data without the need of measuring or monitoring directly each transition. When compared with the direct observation of a transition, the data employed within our approach is in a sense less informative. As with any inference problem, this comes at the price of the need of larger amount of data in order to perform statistically reliable estimates. We have shown that for a reasonable number of lineages (*M* ≤ 3), inference can be performed with a number of data-points which is experimentally reasonable even with a flat prior on the parameters. When increasing the number of populations the number of data becomes significantly larger and might not be fully compatible with experimental needs, therefore potentially requiring to complement information with a non flat prior. Note that an alternative to impose a non flat prior is that to match the moments of the process from the data. Within the BGW formalism this can be done by equating analytically derived moments with empirical values. Such procedure sets a number of additional constraints on the parameters, thereby reducing the amount of data required for inference.

The proposed BGW branching processes involve essentially discrete states and discrete time. While more complicated phenotypic descriptions having continuous states cannot be described within this formulation, one advantageous aspect of the approach presented here is that stochasticity is taken into account in a very natural way. This means that extinction of clones is possible in the model even within clones that are growing on average. This feature cannot be considered directly in continuous models. A valid method to infer properties of phenotypic transitions was proposed in ref.^44^, followed by a number of many successful applications to experimental clonal analysis. Such formulation rather than being based on inference is relying on the process of fitting asymptotic size distribution through the dynamics reconstructed via a master equation. Typically, clonal dynamics is resolved by assuming one unique duplication time for both symmetric and asymmetric cell division, and the population model is postulated *a priori*. Within these limits, asymptotics presented in ref.^44^ might be more reliable than the approach presented here for long observation times and large clones. Our approach is indeed based on a higher degree of synchrony and has quite the opposite behavior, with large asynchronous clones being more difficult to characterize. It is worth noting however that our modeling approach does not allow non-biological rapid cell doublings as would a Poisson distributed doubling time.

One piece of information that is required to apply our approach in the case of multi-lineage dynamics is the number of defined types that compose the population. While this is strictly necessary in the examples presented within the paper, we note that model selection approaches with varying number of populations could be applied in our proposed framework in order to identify the most adherent model to the presented data.

We note that the formally correct way to consider birth-death processes with biologically realistic doubling time distribution and dynamics would be to choose a Bellman-Harris stochastic process rather than a Bienaymé-Galton-Watson. This however comes with significant formal complications and will be left for future developments. It is indeed important to note that our MCEM method is not necessarily bound to BGW processes in particular and its use can be extended to other discrete branching processes or other theoretical models quite simply.

Another aspect to be discussed is the possibility to include cell fate correlations between sister cells, or the possibility of having one progenitor that can follow different types of symmetric and asymmetric cell divisions^13,17,31–33^. The model introduced here includes the possibility for a single progenitor type to transition into any other states through symmetric or asymmetric cell divisions and also includes the possibility of transitions not explicitly linked to cell cycle. Such wide range of possibilities allows our approach to uncover previously unknown modes of tissue homeostasis and growth paradigms. As a side comment, throughout the experimental data considered here, cell death of both siblings (either synchronous or not) was anecdotally observed. Such scenario, which might appear difficult to consider in our approach, could indeed be included by introducing a new cell type which can be reached by symmetric cell division and can only die with probability 1.

In conclusion, we presented a suite of inference methods to identify lineage transitions within a cell population. Our strategy is useful to dissect population hierarchies in growing and treated cell populations and gives access to dynamical features without requiring cell tracking but only cell counts and types. Future research will include the generalization of our approach to other types of branching processes in order to extend the applicability to an even more general set of biological problems.

### Limitations of the study

Discrete stochastic branching processes like BGW do not have clear notion of time and everything is contained within the concept of generation. When putting together inferred probabilities and trying to express these as rates this needs to be taken into account. Also, whenever dealing with experimental data, a number of pre-processing steps needs to be performed to align the data series and estimate the generation time.

As written throughout the manuscript our method works well for progenies that remain substantially correlated for several generations, while it is not suitable *a priori* for Poissonian or similar statistics. Indeed, as a general consideration, BGW branching processes are not suitable to describe large well mixed cell populations as these would show completely asynchronous cell divisions. It is worth noting that in our simulations (SUPPLEMENTARY MATERIAL, sec. 6), estimates based on inference with BGW on realistic data deviates from empirical values of around 10% which is reasonable in the logic of guiding experimental validation.

As detailed above, Bayesian methods rely on writing the full likelihood of a given observation and for many generations (or many observations or many cells). This translates into extremely long polynomials which are hardly manageable with currently available computing possibilities. Such situations can however easily be addressed with the MCEM method which does not need explicit likelihood knowledge. A feasible approach (left for future extensions of the current approach) is to resort to truncated likelihoods.

We note also that BGW are normally spaceless models, where the notion of neighbors is in principle not included. In cases where spatial interactions are necessary this model would not be a natural choice.

## METHODS

### Inference of cell transitions in a BGW branching process

We begin by defining what kind of data we aim at inferring from. As previously described the observations 𝒲 can be thought in different experimentally relevant situations:

1. When tracking is available, we imagine to have access to cell identities and transitions at all times. In general, this means that 𝒲 contains for each cell its lineage, its predecessor and the transitions that each cell performed at the end of each generation. This is a matrix that for each generation contains a variable number of entries (the cells at each generation). As explained in the METHODS section, the properties of the BGW processes are such that the probability of observing a particular configuration does not depend on the particular tree that ends up in that configuration but rather from the number of different transitions observed. Therefore, in practice, here consists of a vector 𝒲 = {*w*_1_, …, *w*_*T*_} of integer numbers, each counting the transitions of each type.
2. In the case of consecutive measurements, the data consists in a set of matrices **Y**, one for each experimental replicate. Each entry in the matrix *Y*_*mk*_ of size *M* × (*n* + 1) represents the number of cells of phenotype *m at* a given generation *k*. Here by *n we* indicate the maximum number of generations followed in the data (*k ∈* {*0*, …, *n*}). It is worth noting that each matrix represents the offspring of one or more cells but all cells are related by genealogical relations in a given matrix. Here 𝒲 = {**Y**^1^, …, **Y**^*R*^} *wh*ere *R is* the number of replicates or datasets collected.
3. In connected end-points measurements each matrix **Y** is an *M ×* 2 matrix where *Y*_*m*1_ contains the zero-th generation and *Y*_*m*2_ contains the *n*-th generation. Here 𝒲 = *{***Y**^1^, …, **Y**^*R*^*}*.
4. In unconnected end-point measurements, each observation is represented by a collection of *M* × 2 matrices, each containing generations 0, *k* for different values of k (for example k ∈{0, …, *n*}) but there is no genealogical relation between cells at different generations, so that effectively one deals with several connected end-point measurements obtained at different unrelated generations. Here 𝒲 = *{***Y**^1^(0), …, **Y**^R^(*n*)*}*

### Modeling cell lineages as stochastic branching processes

In order to be able to infer transitional probabilities in a multi-lineage biological setting we frame our problem into the context of multi-type BGW branching processes^45^. We consider *M* distinct types (or lineages), where individuals (or cells) of the same type are assumed to behave identically, while cells of different types may exhibit distinct behavior. Each cell with lineage *m ca*n generate a number of descendants (daughter cells) between 0 and 2, and the cell type of each daughter cell belongs to one of the *M* possible lineages, while descendants of a cell of lineage *m ca*n in general belong to two distinct lineages *m*^*′*^ and *m*^*′′*^ (either coincident as in the case of symmetric cell divisions or not as in the case of asymmetric cell divisions, see fig. 1A). In terms of BGW formalisms, this translates into a random variable describing the offspring of each lineage *m* at the *n*-th generation that we indicate with 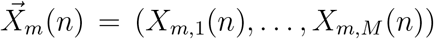. The components 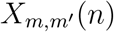 represent the number of descendants of type *m*^*′*^ generated by type *m at* the *n*-th generation. For example a realization 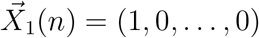 corresponds to a transition 1 *→* 1, where a cell of type 1 was inactive at that generation, 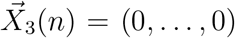 corresponds to 3 → ∅, where a cell of type 3 died, and 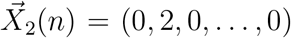 corresponds to 2 → (2, 2), i.e. a cell of type 2 performing cell division.

The population size at the (*n* + 1)-th generation can be expressed in terms of the number *Y*_*m*_*(n*) of cells of each type *m* at the *n*-th generation:

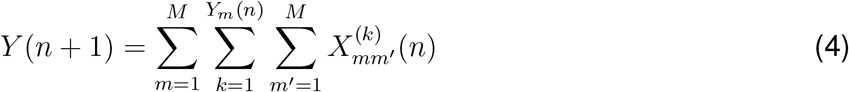

where the index *k* runs on all the *Y*_*m*_*(n*) individuals of type *m* at the generation *n*. Note that 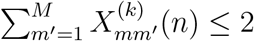.

In the framework of multi-type BGW model, for each type *m*, we can write the probabilities 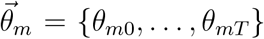 associated to each possible transition that an individual of cell type *m* can perform, where *T* is the total number of different offspring configurations as shown in fig. 1A. It is worth remembering that the ordering of the transitions is arbitrary in the *M⨯T* matrix **Θ** containing all the probabilities *θ*_*mj*_. The total number of transitions *T* for each phenotype *m* can be calculated as follows: the transitions shown in fig. 1A, I-III (i.e. cell death, inactivity and selfrenewal) will be 3, then there will be 3(*M −* 1) for those shown in fig. 1A, IV-VI and (*M −* 2)(*M −* 1)/2 for those in fig. 1A, VII. This gives *T* = (*M* + 1)(*M* + 2)/2.

Each cell type is assumed to act independently and so we have a normalization condition 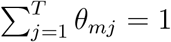 on each parameters 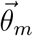 governing the cell behavior, for a total of *M* constraints. The probability generating function (PGF) of the vectors 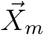 is defined as :

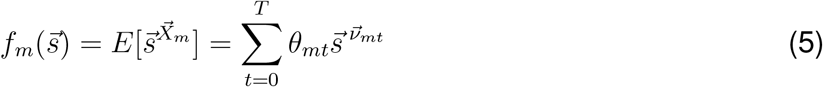

where *s*_*m*_ are the auxiliary variables relative to each phenotype and are grouped in a vector 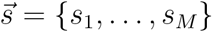, and where we have used the following compact notation 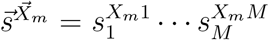. The vector of exponents 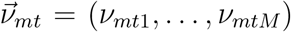 contains the number of cells for each type that emerged from the transition *t*.

Given that the maximum number of offspring is 2, eq.(5) is a polynomial with at most degree 2. The PGFs of the offspring distribution 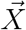 can be used to calculate the population distribu-tions 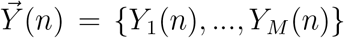 containing the number of individuals for each type at the *n*-th generation by recursive composition. By taking into account the properties of the PGFs, the probability to obtain a specific number of individuals for each type {*Y*_1_(*n*) = *y*_*1*_, …, *Y*_*M*_ (*n*) = *y*_*M*_}, while starting with one individual of type *m* can be written as:

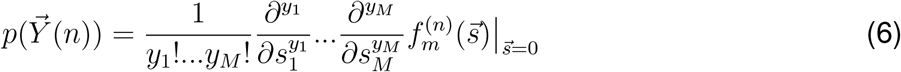

where 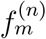 is the recursive composition performed *n* times on *f*_*m*_, i.e.:

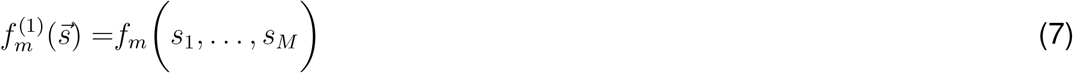

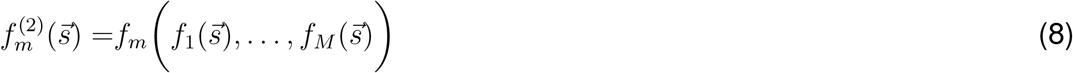

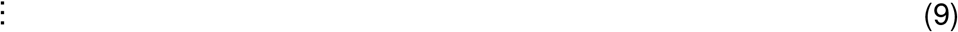

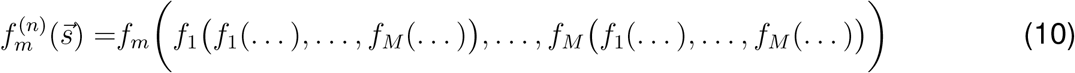

It is important to note that eq.(6) contains all possible trees, or combination of events, that connect a given initial configuration with one cell of phenotype *m* to the generic configuration 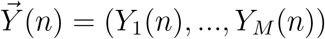. A worked out example for 2 types is given in section 1 of the SI and in fig. S1.

The BGW model just introduced has obvious analogies with the context of cell phenotypes, where each cell type might be thought as a distinct lineage, characterized by a relatively stable transcriptional status. Transitions in this context identify all possible actions that each lineage can undergo. Cell death, inactivity and self-renewal for a given phenotype *m* are indicated in fig. 1A (I-III) and correspond to terms in 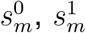 and 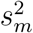 respectively in 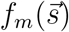. Note that a non-zero probability of inactivity for a given phenotype, while not changing the number of cells in the population, allows some degree of dynamic desynchronization that naturally appears in biological setups. A cell of a given phenotype *m* can also switch to a phenotype *m*^*′*^ as shown in fig. 1A (IV) to take into account extrinsic de/differentiations and would appear as terms in 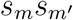 in the corresponding PGF 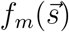. Cell divisions other than self-renewal, can either be symmetric (fig. 1A (V)) or asymmetric (fig. 1A (VI,VII)) where either one or none of the daughter cells is equal to the mother cell and these two terms would be of the form 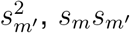 or 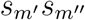 in the PGF. Average growth rates and moments of each lineage can be easily obtained from the PGF^45^. In the proposed model the number of parameters *M·T* grows rapidly with the number of phenotypes with law *M·T* = *M*(*M* + 1)(*M* + 2)/2. Therefore, multi-lineage complicated population structures map into a relatively large set of interpretable parameters.

It is worth noting that if the parameters are all constant, lineage plasticity would be described by our model as just a random event with assigned probability. However, parameters can also be made a function of the number of cells in one or more types, making this model more suitable to describe interaction among cell types or secreted molecules^30^. In this situation, in order to maintain markovianity of the process, one needs to have as many sets of parameters as the number of conditions or number of cells in a given phenotype.

In what follows we show how to implement inference with such a model, by starting from observed data. It is the data in our approach that will select which transitions are compatible with observations and which not, determining the underlying hierarchical structure of the population. Including all possible transitions in the model means that inference is performed even with no prior knowledge on population dynamics. We developed 3 different approaches which give access to different levels of information: analytical inference, extended Gibbs sampler and Monte Carlo Expectation Maximization. Analytical inference is strictly Bayesian, while the Extended Gibbs Sampler (EGS) is a numerical method. Both these approaches output the full probability distribution functions for the parameters (exact and approximated respectively) in situations where we can computationally afford to write the likelihood (small amount of data/low number of cells or sparse data). A third method is presented (the Monte Carlo Expectation Maximization, MCEM) which outputs only point estimates for the parameters and is suited to cases where the amount of data is high and/or the data is sparse, i.e. we do not necessarily have information about intermediate population sizes.

### Analytical Bayesian Inference on BGW branching processes

Bayesian inference applied to BGW branching processes allows to infer the posterior distribution for the model parameters starting from the empirical observation of single or multiple trees^46,47,67–69^. What follows is written as referring to consecutive observations without tracking but can be extended to any of the observations 𝒲 presented in the previous section. We use the letter **Y** to indicate the whole data in 𝒲. Note that while here we derive the approach on the entire dataset at once, one can apply the following formalism for one instance of **Y** at a time, and update the prior at each iteration with the posterior. This approach is followed in the manuscript only to explicitly show the convergence process while adding data but is not computationally efficient.

For now the set of parameters **Θ** is thought as being the same across all datasets which are essentially different observations or realizations of the underlying BGW process. In this case we can start from Bayes’ rule:

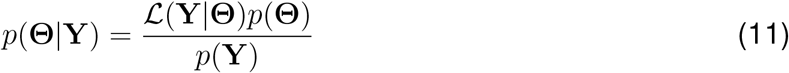

where we are relating the probability distribution of the parameters given the observed data *p*(**Θ**|**Y**) with the prior belief on parameters *p*(**Θ**) and the probability of observing the specific data given a certain value of the parameters (or likelihood) ℒ (**Y**|**Θ**). The latter, within our model, is found through the PGF. We can indeed decompose the likelihood using two distinct assumptions: independence and markovianity. Different experiments/observations are assumed to be statistically independent, and the number of cells at the (*n* + 1)-th generation depends only on the number of cells at the *n*-th generation. In this way we can write the likelihood as a product of the probabilities of each independent observation, which can be easily derived from eq. (6). The term *p*(**Θ**) on the numerator of the r.h.s. of eq. (11) incorporates our belief on the parameters. Among the possible choices we assume a flat prior. This assumption can however be updated with a different and more stringent belief with the immediate effect of decreasing the number of free parameters in eq. (11).

The denominator on the r.h.s. of eq. (11) is known as the marginal likelihood (or evidence) and can be written as

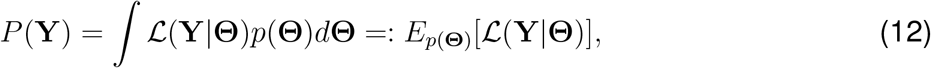

where the integration is taken over the whole parameter space, taking into account the normalization condition. Eq. (11) leads us to the posterior *p*(**Θ**|**Y**), which is the probability distribution of the parameters of our BGW model given the observed data. This is a multivariate function which needs to be marginalized to find the probability distribution *p*(*θ*_*mj*_|**Y**) for each single parameter. Point estimators like the mean and the variance of the distribution can be easily calculated starting from the marginal distribution above.

The analytical approach here presented can be employed in several experimentally relevant situations, with a number of limitations mostly originating from computational costs. A first interesting case is that of cell tracking: in the logic of being able to track all cell transitions the likelihood for each observation is essentially that of a particular tree, and the likelihood of all independent observations just the product of all likelihoods. Inferring parameters in this situation is therefore conceptually simple as applying eq. (11) as is. Another interesting case would be that of having the counts for each type, or the total number of cells at each generation. In this case one would construct a likelihood with the conditioned probabilities and again apply directly Bayes’ formula. Finally, this approach could also be applied when one only has information regarding the count of individuals of each type on non-consecutive generations, a situation typically encountered when dealing with fixed samples at a given time. In this case the likelihood would contain all possible trees to reach a given configuration. A plot of an example of a full posterior is shown in fig. S10. The numerical implementation of the presented approach is described in the METHODS and a sketch is given in fig. S9A.

One severe limitation of the proposed analytical inference is that computational cost becomes rapidly unpractical when increasing either the number of data, the number of cells or the number of lineages and hence the number of parameters. In such situations the likelihood becomes a significantly long list of monomials that while trivial to integrate constitute a computationally prohibitive challenge on standard computers. Analogously, if the data only contains information on a late number of generations, the number of trees to satisfy the observation can grow quite rapidly and become computationally intractable. Therefore, alternative methods are needed, based on numerical simulations that possibly avoid the explicit calculation of the marginal likelihood.

### Approximate Bayesian Inference with the extended Gibbs sampler

The explicit knowledge of the analytical likelihood allows to extend previous work^69^, belonging to the class of Monte-Carlo Markov-Chain algorithm, to the more general case of inferring the parameters of a multi-type BGW process without the need of generation-contiguous observations. The approach developed here, can be applied to any dataset 𝒲 with the exception of the case with tracking which has no reason to be tackled with this method (see ref.^67^ for an extension towards non-parametric distributions). We use the letter **Y** to indicate the data in 𝒲 (without tracking).

As explained at the end of the previous section, a problem of the analytical setup is that when the number of cells is large, for any given dataset 𝒲 (without tracking information), there are multiple trees that end up in the same configuration, resulting in a likelihood which is the sum of several terms each corresponding to one of these trees. Each of these trees can be indicated by a matrix **Z** containing the count of all the possible *Z*_*mt*_ transitions for each phenotype *m*, where *m* = 1, …, *M* and *t* = 1, …, *T*. The exact knowledge of the tree **Z** determining the configuration **Y** with a given initial condition requires tracking each cell at all times. However **Z** can also be considered a latent variable, and this assumption still allows to express the likelihood:

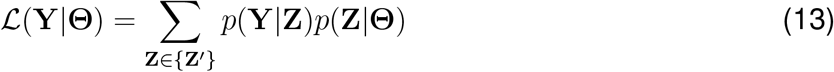

where the summation is over the set of possible trees {*Z*^*′*^} compatible with the observation **Y**. The idea behind the Gibbs Sampler^69^ is to sample directly from the posterior distribution, knowing that it is proportional to the likelihood via the prior:

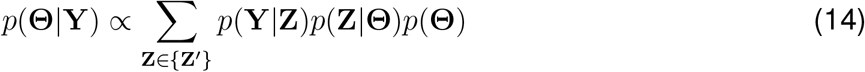

To sample from the posterior distribution one can sample a first parameter matrix **Θ**^*∗*^ drawing from the prior distribution. If all types are independent the prior can be factorized into marginal priors over each phenotype, and each marginal 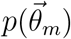 can be expressed in terms of a Dirichlet distribution:

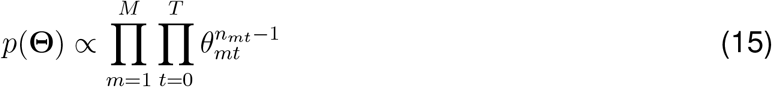

where *n*_*mt*_ reflects the shape of the chosen prior distribution.

At this point, without tracking information, one needs to sample among all the possible trees that are compatible with the observation **Y**, which is formally done in eq. (14). Such information however is contained within the analytical likelihood, which lists all the different trees compatible with the observation as a sum of terms. Therefore the likelihood itself can be used to sample among all the trees as follows. Each monomial of the likelihood corresponds to a possible tree **Z**. We can therefore construct a vector by substituting the values **Θ**^*∗*^ into each monomial of the likelihood. Upon normalization, such a vector gives the actual probability of occurrence of each tree *p*(**Z**|**Θ**) and can be used to sample trees. At this point it is sufficient to exploit the fact that a Dirichlet distribution and the product of multinomial distributions are conjugated to find an instance of the posterior:

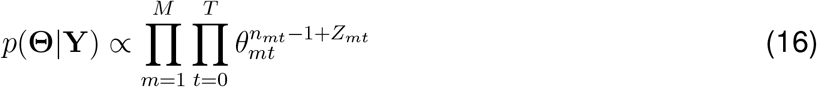

This posterior distribution can finally be used to sample a new set of values **Θ**^*∗*^.

The iterative use of this procedure produces a sequence of values {**Θ**^*∗*^} which distribution is known to converge to the true marginal distribution. A sketch of this approach is presented in fig. S9B, and the comparison with the analytical method is presented in fig. S11. It is worth noting that this approach generalizes that presented in ref.^69^, which only allows to infer from data series containing generation-contiguous observations, i.e. *Y* (1), …, *Y* (*n*).

While the proposed version outperforms the analytical methods in a number of situations, it still relies on the ability of writing and calculating an exact likelihood function, which can be prohibitive in a similar set of cases as the previous method, making the use of this method suitable for intermediate number of cells and observations. Both the analytical method and the extended Gibbs sampler (EGS) suffer computational regimes that increase the exponents in the likelihoods (i.e. a higher number of cells and a higher number of observations) or those that increase the number of terms in the likelihood such as incomplete or non contiguous observations. A phase portrait containing these limitations is shown in fig. 1C. Such considerations call for yet another approach with better computational performances.

### Inferring maximum likelihood estimators with Monte Carlo Expectation Maximization

In this section we limit our attention to connected non-consecutive observations (indicated as usual with **Y** which represents data in 𝒲). In this scenario, the likelihood contains a possibly very large number of possible trees or forests that are all compatible with the observed data. Given these challenges, it is computationally very difficult to compute the posterior distribution for the parameters with the previously described approaches. Therefore, we developed a Monte Carlo Expectation-Maximization algorithm (MCEM) to efficiently estimate the parameters by maximizing the log-likelihood. If *y* is one particular observation within the set **Y**, the set of parameters 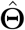 maximizing the log-likelihood is:

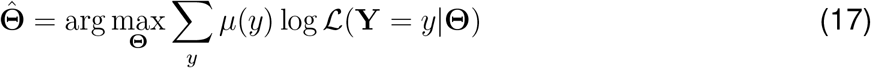

where *μ* is the empirical distribution:

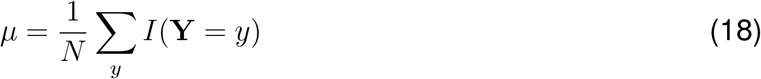

and I is an indicator function.

As with the EGS, we write everything in terms of the hidden variables **Z**, containing all possible ways to go from an initial population to a given population at a future generation:

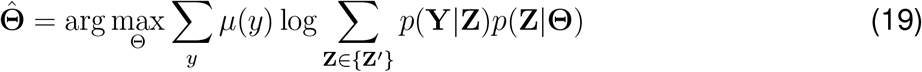

This problem does not have a closed form solution. However the MCEM algorithm can be used to maximize the likelihood iteratively, by means of a surrogate function that possesses the same extrema of the likelihood. A surrogate function ℬ (**Θ**) can be found exploiting Jensen’s inequality: log(*E*_*p(x)*_[*f(x*)]) *≥ E*_*p(x)*_[*log f*(*x)*] and can be written as:

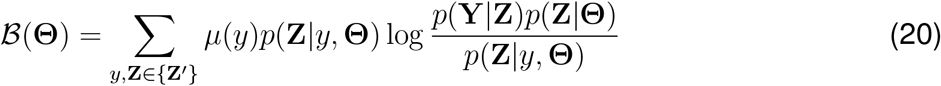

where the terms 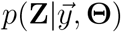 are usually called responsibilities and can be written as 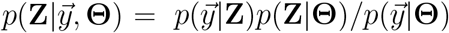.

In our multi-type BGW the term *p*(**Z**|**Θ**) can be written as:

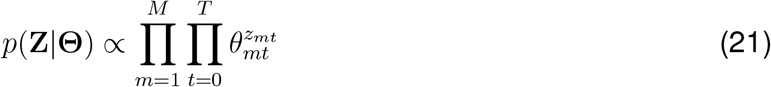

The maximization of ℬ(**Θ**) can be obtained with the method of Lagrangian multipliers with *M* constrains. This procedure lead us to maximum-likelihood estimation for each parameter

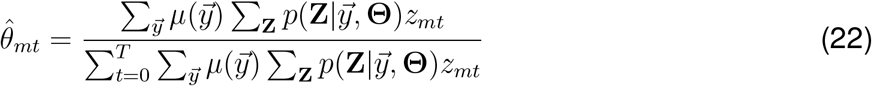

The terms in the numerator required for the conditional probabilities can be theoretically calculated optimizing the surrogate function ℬ(**Θ**) with respect to the responsibilities. However, it is possible to overcome this procedure by estimating the numerator by means of more efficient numerical methods. The previous numerator is approximated by the quantity:

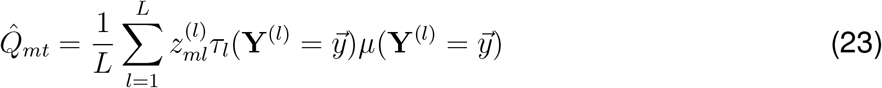

where the average is performed over *L* different numerical simulations and 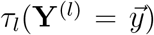 represents inter-arrival time between two identical observations defined mathematically as

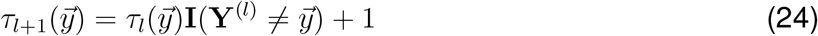

with initial condition 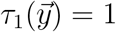. With this estimators, each parameter *θ*_*mt*_ can be estimated through the formula

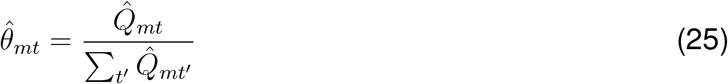

This numerical approach is computationally effective even for relatively large number of cells and parameters and covers a wide range of experimentally relevant situations as shown in the next sections. Such efficiency however comes at the price of sacrificing the entire probability distribution function in favor of point estimators 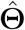. A graphical sketch for this method is shown in fig. S9C. Also, note that the method proposed here can be easily extended to the case of different initial cell numbers or types. The extended version is discussed in the SUPPLEMENTARY MATERIAL, sec. 7.

### Implementation of analytical Bayesian inference with Mathematica

In order to perform analytical Bayesian inference we made use of the software Wolframe Mathematica 13.2. Given an observation **Y** we first generate the likelihood ℒ (**Y**|**Θ**) using the PGF of the model considered. The aim is then to calculate the posterior using Bayes’ formula:

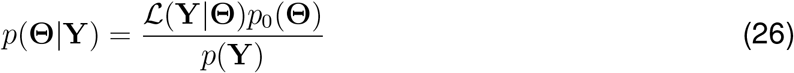

where the prior *p*_0_ is taken to be flat, i.e. constant. The non trivial step here is to calculate the term in the denominator, *p*(**Y**), which can be written as

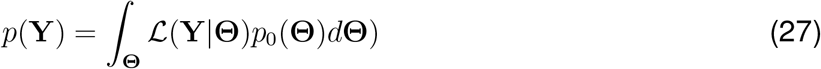

where the integration is performed over the entire parameter space and takes into account all the *M* constrains. For the simplest case of one and only phenotype *M* = 1 this formula reduces to:

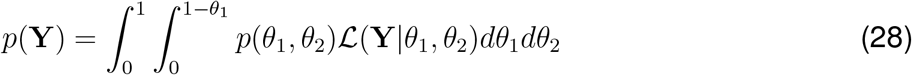

where the integration is performed over only two parameters thanks to the constraint *θ*_0_ = 1 *− θ*_1_ *− θ*_2_.

It is worth noting that the likelihood for a general BGW process contains only polynomial terms and is therefore theoretically straightforward to integrate analytically. However, when the number of observations becomes large or there are many possible trees that explain one given observation, it can contain a significantly large number of terms and represents therefore one of the most computationally intensive tasks when implementing eq.(26).

Note that this approach can be taken iteratively or in one step. When computing iteratively one basically repeats the calculation in eq. (26) for each observation separately using the posterior at step *n* as the prior for step *n* + 1. Alternatively one can calculate the full likelihood of all the observations at once and perform a one step calculation.

Once the posterior is obtained we can calculate the marginal distributions by integrating over all the parameters but one. The moments of the marginal distributions are also readily calculated starting from the marginal PDF. Two sample Mathematica scripts covering the cases of one type and two types can be found on our dedicated github: https://github.com/AndreaPiras21/Inference-Cells/.

### Implementation of numerical inference: the extended Gibbs sampler

The logic of a Gibbs sampler algorithm is to iteratively draw values from fully conditional distributions. In our setting, the fully conditional distribution 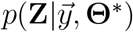 is provided directly by the analytical likelihood. Indeed, we can draw a particular set of trees by exploiting the fact that the likelihood contains all possible admissible sets of trees linking two separate generations that agree with the initial condition and the final observation. The choice of the tree set depends on the current parameter values 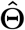, which are initially drawn from a flat prior and subsequently drawn from a set of Dirichlet distributions, –one for each phenotype– updated based on the previously selected set of trees, 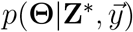.

The obtained series of drawn 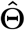 builds our approximated numerical posterior distribution, and each value 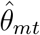 can be exploited to build the approximated marginal distribution of the transition *j* with respect to phenotype *m*. A numerical implementation of this method in Mathematica can be found on our dedicated github: https://github.com/AndreaPiras21/Inference-Cells/.

### Implementation of numerical inference: Monte Carlo Expectation Maximization

The Monte Carlo Expectation-Maximization (MCEM) algorithm estimates transition probabilities in a Bienaymé–Galton–Watson (BGW) process by simulating cell lineage trees. These trees, which represent the full developmental history of a cell population, are typically not observed in experiments and thus constitute latent variables. However, they can be relatively easily simulated using a BGW process. Starting from an initial set of parameter values 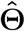, the algorithm simulates multiple cell lineage trees. For each simulation *l, it rec*ords how many times a specific transition *t* occurs for a given phenotype *m*. At the same time, it also tracks the number of steps elapsed between two identical realizations (eq. (24)). The inverse of the mean of this quantity becomes essentially the probability of observing that particular realization. This inferred probability is then combined with the empirical frequency of the realization in the observed dataset. Together, these quantities are used to update the parameter estimates 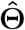. The update is performed through the empirical mean 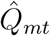 in eq. (23) which is iteratively updated while samples are generated using the following update rule:

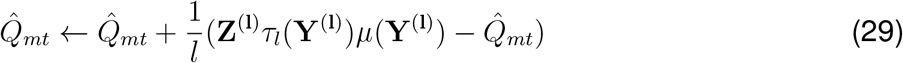

Finally, the updated parameters are obtained using eq. (25), and the entire process is iterated until the estimates converge to a local maximum. A Python script implementing MCEM for one type BGW process can be found on our dedicated github: https://github.com/AndreaPiras21/Inference-Cells/.

### Numerical simulations of BGW processes

Simulations of the BGW process are generated using a Python script. Given an initial number of cells with a known phenotype composition, we simulate a given number of generations. At each generation, for each cell of phenotype *m*, we randomly select a possible transition according to the offspring distribution associated with that phenotype. This is simply done by throwing a random number and deciding which transition within 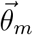 the cell will undergo. We then update the number of cells for each phenotype, taking into account that different transitions lead to different changes in the sub-populations. In our setup, the only possible changes for a sub-population due to a cell of phenotype *j* are: Δ*n*_*j*_ ∈ {1, 0, 1, 2}. It is important to note that the order used to list the transitions is arbitrary, and alternative orderings are equally valid. Finally, when feedback mechanisms are incorporated, the offspring distribution is updated after each change in the sub-populations, according to a predefined mathematical rule. This allows the model to capture scenarios in which the parameters depend on the total number of cells in the population or on the number of cells within specific sub-populations. Python scripts to simulate BGW processes can be found on our dedicated github: https://github.com/AndreaPiras21/Inference-Cells/.

### Experimental procedures, materials and methods

MG-63 cell line was purchased from Interlab Cell Line Collection (ICLC). The cell line was kept in stock in our institute’s cell culture facility, which re-authenticates cells by applying the PowerPlex 16 HS System (#DC2101, Promega, USA) and tests for Mycoplasma contamination with a PCR Mycoplasma Detection kit (#G238, Applied Biological Materials Inc., Canada).

MG-63 were cultured at 37C under 5% CO2 humidified atmosphere in DMEM low glucose (#ECM0070L, Euroclone S.p.A., Milano, Italy). Culture medium was supplemented with 10% fetal bovine serum (#A5256701, Gibco BRL, ThermoFisher, Waltham, MA, USA), 100U/mL of penicillin and 100*μ*g/mL streptomycin (#ECB3001D, Euroclone S.p.A., Milano, Italy) and 2mM L-Glutamine (#ECB3000D-20, Euroclone S.p.A., Milano, Italy).

Cells were cultured in standard tissue culture plastic dishes (Falcon) and imaging dedicated supports (#80826 *μ*-Slide 8 Well ibiTreat; #89626 - *μ*-Plate 96 Well Square, ibiTreat; IBIDI, GmbH, Grä felfing, Germany). MG-63 cells transduced with lentiviral vectors indicated below were cul-tured at low density (125/well-96 Well Square or 500/well-8 well) on day 0, in 300 *μ*L of complete growth medium. The next day, we performed live imaging choosing only single-cell-fields, with an inverted wide-field microscope Nikon Ti2 (Nikon Instruments, USA) with a 20X 0.75 NA objective and a digital camera (IRIS 15; Photometrics, USA) and an incubator to keep the plate stably at 37C and 5% CO2. Images were taken every 20 minutes and the time-lapse assay lasted 96 hours.

Plasmids were purchased from Addgene. pLentiPGK DEST H2B-iRFP670 was a gift from Markus Covert (Addgene plasmid # 90237^70^). pBOB-EF1-FastFUCCI-Puro was a gift from Kevin Brindle & Duncan Jodrell (Addgene plasmid # 86849^65^). Cells were segmented with the help of Ilastik^71^, Cellpose^72^ and custom scripts written in Matlab (The Mathworks). Briefly, timelapse sequences of MG63-H2BiRFP670 were segmented with Ilastik and tracked by means of a custom script written in Matlab. Obtained tracks were manually inspected for occasional mistakes. By this means we could either obtain all the transitions 1 observed in each data series or compute the total number of cells at all times and process data 1 as specified in the next section. 1

MG63-FastFUCCI timelapse sequences were segmented by creating a linear combination of red 1 and green channel and applying Cellpose. Then cells were tracked as before and tracks were 1 manually corrected. Red and green signals corresponding to segmented cells were rescaled to 1 avoid time dependent fading (which we found relevant particularly for the green channel). With 1 the obtained point distribution, and in order to define the state (green or red) of each cell, we built a Gaussian Mixture Model with two components for each data series. Cells were then assigned to either class at all timepoints. In order to avoid wrong classification we isolated cells that were not clearly assigned to either class (by filtering for the value of the posterior probability of assignment to either class). We found that these would correspond to ‘phenotypic switches’, i.e. either cells that just divided, where red intensity was not strong enough yet, or cells that were ceasing to express red to start expressing green. In such cases we assigned the closest future state. Tracks for each data series were either transformed into counting of all transitions or processed as explained hereafter (fig. S12E,F).

## Experimental data processing

In order to perform inference on experimental data we first determined the average doubling time *T*div for a given set of experiments and then constructed sampling times spaced away by a cell doubling time. Experimental data (once processed and segmented) were expressed as temporal sequences of number of cells *n*(*t*) where the frame rate is either 10 or 20 minutes. In order to find a set of sampling points to convert the finely resolved experimental data series into BGW branching process-like data series, we identified, in each data series, all regions with 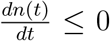 Once excluded connected regions shorter than 4 hours, we identified *n*_*e*_ regions (typically 4 or 5 for each data series), corresponding to generations. For each of these regions we identified the first and last time-point: [*t*_2ℓ*−*1_, *t*_2ℓ_], with ℓ = 1, …, *n*_*e*_ and the midpoints: 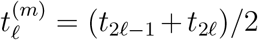. The corresponding *n*_*e*_ sampling points 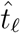 were found by minimizing the function:

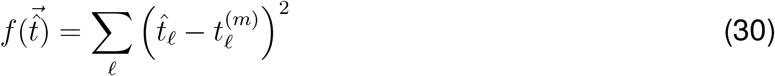

over 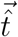, with the constraints:

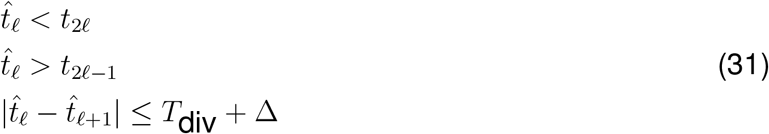

where *T*_div_ is the average doubling time over trajectories measured for each cell and Δ is a tolerance set to *T*_div_/2. Such a system of constrained nonlinear equations was minimized with Matlab (see fig. S12C). Data series with misidentified plateau were manually corrected.

For MG63-Fast-FUCCI we sampled every half average doubling time in order to accomodate for the cell cycle transitions from G1 to S/G_2_/M. We used the same nonlinear programming solver in Matlab to impose sampling points at the midpoint of each plateau which was identified by considering regions where the number of red (or green) cells was equal to the total number of cells. Each sampling time was again picked to fall within a plateau and to be a half duplication time away from the following and previous sampling times.

Cells trajectories for the case of contact inhibition were only aligned on the first cell division.

## RESOURCE AVAILABILITY

### Lead contact

Requests for further information and resources should be directed to and will be fulfilled by the lead contact, Alberto Puliafito.

### Materials availability

This study did not generate new materials.

### Data and code availability

Instances of the codes are available on GitHub at the following link:

https://github.com/AndreaPiras21/Inference-Cells/

## ACKNOWLEDGMENTS

This work was supported by AIRC (Associazione Italiana per la Ricerca sul Cancro) via grant MFAG-25040 to APu, IG-23211 to LPr and 5×1000 grant 21091 to AB; FPRC 5×1000 Ministero della Salute 2022 CARESS to APu; MUR (Dipartimenti di Eccellenza DM 11/05/2017 n262) to the Department of Oncology, University of Turin (2023-2027 14586 DIORAMA); Italian Ministry of Health, Ricerca Corrente 2025. We acknowledge fruitful discussions with Livio Trusolino and Sabrina Johanna Fletcher.

## AUTHOR CONTRIBUTIONS

Conceptualization: APi, AC, APu; Data curation: APi, APu; Formal analysis: APi, AC, APu; Funding acquisition: LPr, APu; Investigation: APi, FG; Methodology: APi, AC; Project administration: AC, APu; Resources: LPr, APu; Software: APi, AC; Supervision: AC, APu; Validation: all authors; Visualization: APi, LPi, AC, APu; Writing – original draft: Api, APu; Writing – review & editing: all authors;

## DECLARATION OF GENERATIVE AI AND AI-ASSISTED TECHNOLOGIES IN THE WRITING PROCESS

Statement: During the preparation of this work the author(s) used ChatGPT in order to improve english syntax and to assemble plots with the package Matplotlib for Python. After using this tool/service, all authors reviewed and edited the content as needed and take full responsibility for the content of the published article.

