## Supplementary Material for "Inference of lineage hierarchies, growth and drug response mechanisms in cancer cell populations – without tracking"

**4 MCEM convergence 12**

**5 Implementation of feedback on the total population for multi-lineage models 13**

**6 Inferring from non-clonal or asynchronous experimental data 15**

**7 Monte Carlo Expectation Minimization with variable initial number of cells 18**

**8 Initialization strategies for MCEM inference 19**
- 8.1 Moments of a Multi-type Galton-Watson model . . . 19
- 8.2 Moment Matching . . . 22

**9 Supplementary figures 25**

### 1 Worked-out likelihoods for BGW branching process with 2 cell types

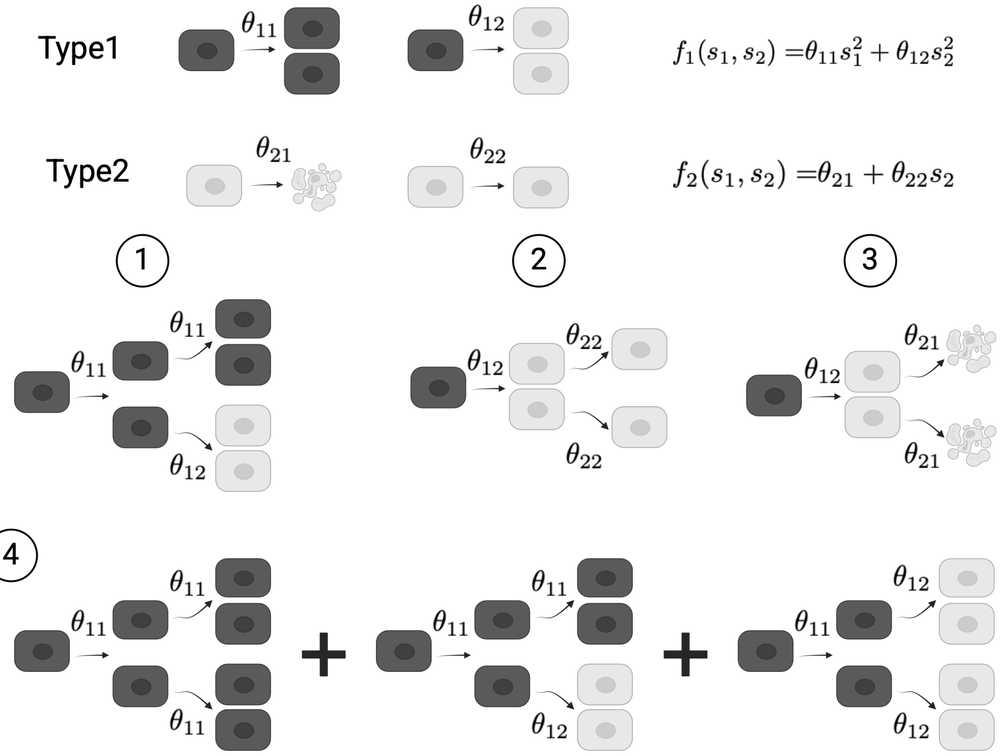

Figure S1: **Calculation of the likelihood for a 2-type BGW process** In this example, admitted transitions for each phenotype (1 and 2) are drawn along with their respective PGFs (top). Circled numbers represent 4 different observations corresponding to different trees at the 2nd generation with a particular number of individuals for each phenotype.

We here explicitly develop the likelihood of observing a set of particular realizations as indicated in fig. S1.
The PGFs in this case can be written as:

$$f_1(s_1, s_2) = \theta_{11}s_1^2 + \theta_{12}s_2^2 \tag{1}$$

$$f_2(s_1, s_2) = \theta_{21} + \theta_{22}s_2^2 \tag{2}$$

In case ① (see fig. S1) we sought to determine the probability of observing one black cell dividing into two black cells which then respectively divide into two black cells and two white cells. This can be computed starting from the PGF as:

$$p(y_1(2) = 2; y_2(2) = 2) = \frac{1}{4}\frac{\partial^2}{\partial s_1^2}\frac{\partial^2}{\partial s_2^2}f_1(f_1(s_1, s_2), f_2(s_1, s_2))|_{\vec{s}=0} = 2\theta_{11}^2\theta_{12} \tag{3}$$

Analogously, case ② represents the tree where one black cell divides into two white cells which then both standby. This can be worked out as:

$$p(y_1(2) = 0; y_2(2) = 2) = \frac{1}{2}\frac{\partial^2}{\partial s_2^2}f_1(f_1(s_1, s_2), f_2(s_1, s_2))|_{\vec{s}=0} =$$

$$= \frac{1}{2}\frac{\partial^2}{\partial s_2^2}[\theta_{11}(\theta_{11}s_1^2 + \theta_{12}s_2^2)^2 + \theta_{12}(\theta_{21} + \theta_{22}s_2)^2]|_{\vec{s}=0} = \theta_{12}\theta_{22}^2 \tag{4}$$

Case ③ corresponds to one black cell dividing into two white cells then both dying:

$$p(y_1(2) = 0; y_2(2) = 0) = f_1(f_1(s_1, s_2), f_2(s_1, s_2))|_{\vec{s}=0} = \theta_{12}\theta_{21}^2 \tag{5}$$

A slightly more complex case is ④ which corresponds to the probability of having 4 cells of any type at generation 2. This last case corresponds to three independent possible trees, whose probabilities add up:

$$\begin{aligned} p(y_1 + y_2 = 4) &= \left(\frac{1}{4!}\frac{\partial^4}{\partial s_1^4} + \frac{1}{4}\frac{\partial^2}{\partial s_1^2}\frac{\partial^2}{\partial s_2^2} + \frac{1}{4!}\frac{\partial^4}{\partial s_2^4}\right) f_1(f_1(s_1, s_2), f_2(s_1, s_2))|_{\vec{s}=0} = \\ &= \theta_{11}^3 + 2\theta_{11}^2\theta_{12} + \theta_{11}\theta_{12}^2 \end{aligned} \tag{6}$$

### 2 Worked-out likelihoods for different data types in a BGW branching process with 1 cell type

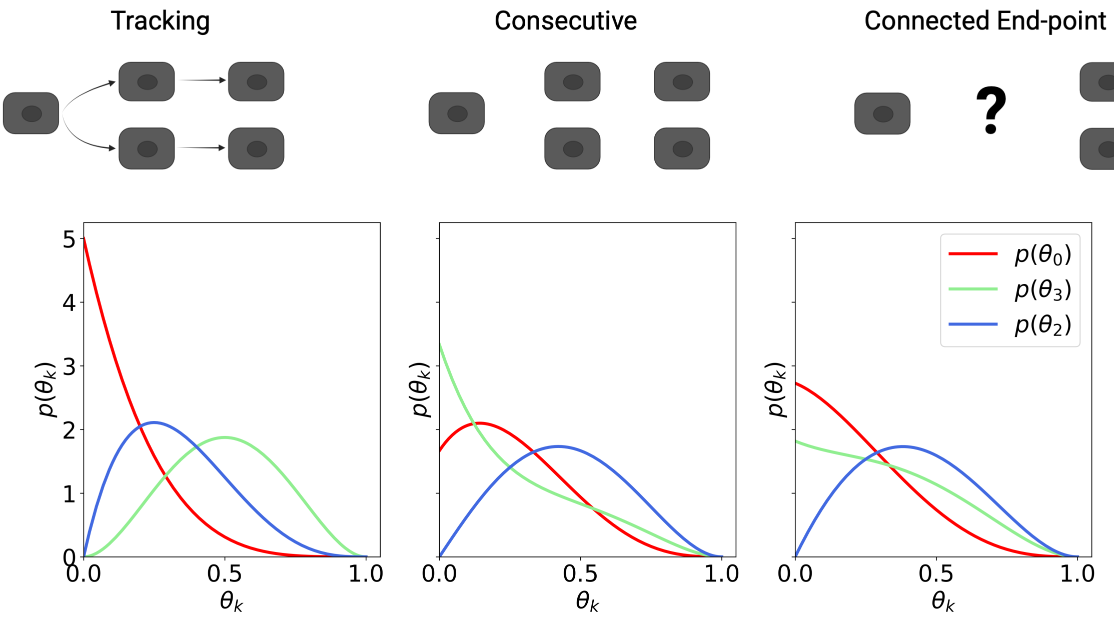

Figure S2: **Analytical inference for three different data-types.** Observations are indicated on top of the figure and the corresponding set of marginal posteriors is drawn below each corresponding sketch.

Here, we explicitly consider an example of inference starting from different types of available data. Suppose we are monitoring two generations of a single-type BGW process with cell death, standby and division indicated by probabilities $\theta_0, \theta_1, \theta_2$ as usual. We consider: i) the full tracking case, where we observe one division at first and then two inactivations, ii) the consecutive measurements case where we observe the number of cells at any generation (here: one, two and two), iii) the case where we only observe the initial and final number of cells.
The tracking case is represented by the following likelihood:

$$\mathcal{L} = \theta_1^2 \theta_2 \tag{7}$$

This corresponds to a single tree where one cell divides first and then the two daughter cells perform a standby. The marginal posteriors correspondingly indicate a larger probability for inactivation and division, while death probability peaks on zero.
The consecutive case can be worked out by multiplying the probabilities of observing each transition ($1 \rightarrow 2$ and $2 \rightarrow 2$), to obtain:

$$\mathcal{L} = \theta_1^2 \theta_2 + 2\theta_0 \theta_2^2 \tag{8}$$

where now there are two possible distinct independent trees one in which there is adivision followed by two standby, and the second where one of the daughter cells divides and the other dies (the factor two follows from the exchangeability of the two cells).
As a final case we consider the case where the configuration at the intermediate generation is not observed. The likelihood can be calculated as previously:

$$\mathcal{L} = \theta_1^2 \theta_2 + 2\theta_0 \theta_2^2 + \theta_1 \theta_2 \tag{9}$$

where we have now a a contribution from a third tree which corresponds to one cell standing by and then dividing.

We remark that in these three cases the same underlying process yields different inference results as shown in the marginals in fig. S2. Note that the maximum of the marginals for parameters $\theta_0$ and $\theta_1$ is inverted when switching from tracking to consecutive measurements.

### 3 Lineage inference in one cell-type BGW models of cell populations

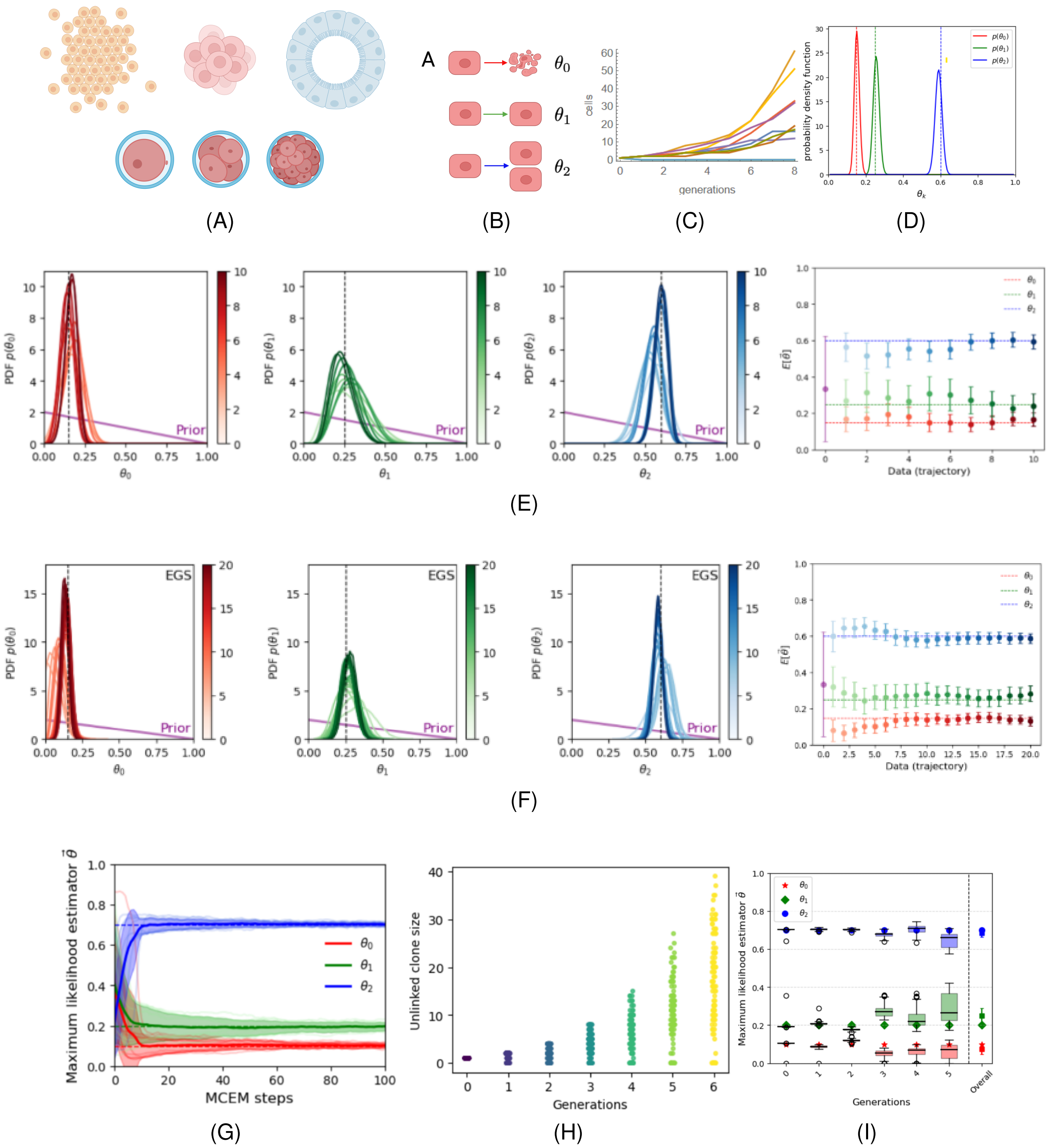

Figure S3: Caption is on the next page.

Here we describe how to apply the different developed methodologies to a number of biologically relevant situations, represented by *in silico* data generated by numerical simulations of BGW branching processes.

Figure S3: **Free expansion of a 1-type cell population** **A** Illustration of biological applications of the 1-phenotype BGW process (top row from left to right: a cell colony, a spheroid or tumoroid, a cyst; bottom row: a developing embryo). **B** Transitions in a 1-type BGW process representing situations as in panel A. True values used were $\vec{\theta}^{\text{true}} = (0.15, 0.25, 0.6)$. **C** Plot of $10$ different realizations of a 1-phenotype BGW branching process. All simulations start with one cell and stop after 8 generations. **D** Marginal distributions obtained with the growth curves shown in C, assuming single cell tracking has been performed. **E** *Left* Analytical marginal posteriors for each iteration are shown in different shades of color and peak more and more as the number of processed data increases. Data are the same as in panel C with no tracking information. *Right* Mean value of the inferred marginals for each parameter during the iterative process, compared to the respective true value (dashed lines). **F** *Left* Marginal posteriors computed with EGS by extending the dataset and considering $10000$ drawing points. *Right* Comparison between the mean of the distributions and the respective true value. **G** Maximum-likelihood estimations of the three parameters performed with the MCEM algorithm. The dataset consists in $1000$ endpoint measurements after 8 generations with no access to intermediate states and true values $\vec{\theta}^{\text{true}} = (0.1, 0.2, 0.7)$. **H** Representation of data collection where each dot is one colony or tumoroid independent from all others. **I** Inference of parameters from data in panel H obtained by using MCEM algorithm. Note that inferred values are consistent with the true values $\vec{\theta}^{\text{true}} = (0.1, 0.2, 0.7)$.

#### 3.1 Free clonal expansion

We consider first the simple case of a single cell type. This example can be linked to several biological contexts (fig. S3A), such as a single cell developing into a cell colony, a spheroid, an tumoroid or a developing embryo in the zygote to morula stage, prior to any cell fate specification. The possible transitions for each cell are cell death, standby and self-renewal, as illustrated in fig. S3B. In order to present the different scenarios where our inference approach can be applied we numerically simulated a BGW branching process and generated different datasets corresponding to different experimental setups all with the same parameter values, as shown in fig. S3B. A plot of the typical set of realizations used is shown in fig. S3C. Note that, at variance with continuous models, the probability of extinction is strictly positive in any nontrivial BGW branching process, and therefore it is possible that a single cell does not give rise to a progeny even if on average the population is growing.

##### 3.1.1 Tracking all cells

First we consider the case where cell tracking is available, i.e. we assume full knowledge of the history of each single cell in terms of transitions on all generations. To this aim we consider ten distinct data sequences generated by numerically simulating a BGW process, starting from a single cell and lasting 8 generations.
This situation can be easily tackled analytically. Across the whole dataset, $107$ cells died, there were $180$ standbys and $415$ cell divisions. The likelihood of such observation can be obtained by factorizing the single tree likelihoods or just considering the total number of events: $\mathcal{L}(\vec{z}|\vec{\theta}) = \theta_0^{107}\theta_1^{180}\theta_2^{415}$. The one final step is to obtain the posterior from such a likelihood which can be easily done by recalling normalization conditions for Dirichlet distributions. A direct application of Bayes' rule with a flat prior and such a likelihood then leads to the posterior shown in fig. S3D. This result is in agreement with the true values used to generate the observations and the width of the distributions expresses the statistical uncertainty originating from the finite number of data points used to infer the parameters.

Thanks to the mathematical properties of BGW processes, cell tracking is phrased into knowing all possible transitions that occurred within a given number of generations rather than really knowing all the different trees. Therefore, in an experimental situation, the information required is the knowledge of how many cells died, how many cell division were observed, while the standbys are automatically calculated by counting the total number of cells, as shown below. This represents an experimentally relevant point as one can use temporally well resolved timelapse experiments and recognize transitions (e.g. mitoses, cell death) without the need to explicitly track cells one by one.

##### 3.1.2 Consecutive measurements with no tracking

Our methodology can also be used when single cell tracking is not available and, for example, we only have access to the total number of cells at each generation, rather than the lineage history of each cell. We therefore consider the same dataset as before but assuming no direct connection between data at different generations. This case was analyzed using our analytical method and the results are shown in figureS3E. Here, we present an iterative approach, where data are incorporated one by one into the likelihood, providing insights about the convergence of the estimations. Our results show that, even without recurring to cell tracking, parameters can be inferred and the distributions are concentrating around the true values.
Analogous conclusions were reached by employing the EGS to a larger dataset, where now the marginals are computed numerically (as shown in fig. S3F). Note that the latter approach is computationally more efficient than the analytical one which is why more sequences, or datasets with more cells can be analyzed.

##### 3.1.3 Connected end-point measurements

We now turn our attention to the case when only the population size at the initial and final generations is measured. This corresponds to an experiment where single cells are seeded and grown for an appropriate time, assuming no mixing is occurring, and where cells are then fixed and counted at the end of the experiment. In this context, 'no mixing' means that cells coming from different ancestors are never exchanged, i.e. they are either very far away or do not move appreciably. This situation can be theoretically addressed both by the analytical and the EGS methods, but becomes rapidly intractable when the number of elapsed generations increases. Such data series can however be analyzed using the MCEM algorithm. This algorithm requires a relatively larger amount of data to produce reliable estimates in contrast with the two previous approaches. To show the power of this approach, we considered a much larger dataset, starting from a single cell, and including only the counts at the 8-th generation. This dataset was then processed with the MCEM algorithm and the convergence to the true values was reached after only a few steps, as shown in fig. S3G.

##### 3.1.4 Unconnected end-point measurements

Finally, we consider a situation in which data from tumoroids or cell colonies are independent at each generation, i.e. each observation is obtained by seeding a single cell and then fixing at a given generation and the experiment is repeated for different final generations, as shown in fig. S3H. Data generated in this way does not allow to the use conditional probability on consecutive generations $p\big(y(n+1)|y(n)big)$ since for a given data-point $y(n+1)$ the corresponding $y(n)$ is unknown.
One possible approach is to marginalize over all possible $y(n)$, giving $p\big(y(n+1)\big) = \sum_y p\big(y(n+$

$1)|y(n))$. Inference with this approach however leads to a larger uncertainty with respect to when the previous data point is known, as it indeed includes fewer transitions and hence less information. Furthermore, this approach becomes computationally intensive for large population sizes or large number of types as the number of possible previous conditions to sum over becomes large. To overcome these issues we approximate the marginalization by summing just over the populations sizes $y$ observed at the previous generation $p(y(n+1)) = \sum_y p(y(n+1)|y(n)) \simeq \sum_{y_{obs}} p(y(n+1)|y(n))$.
In order to perform inference we then apply MCEM by using $100$ observations for each generation, as shown in Fig. S3I. In this way we can recover inferred parameters for each generation of observations and such estimates can be combined to give a more precise global estimate over the whole dataset.
It is worth remarking that this approach does not require cell tracking nor consecutive observations. As such, it is ideally suited to the case when measurements are based on fixed snapshots obtained at different times, a situation which is frequently encountered experimentally. While this is admittedly a less rigorous methodology than the other ones presented in this section, it can however prove useful in experimental setups, provided that data for each generation are sufficiently representative of all possible realizations.

#### 3.2 Clonal expansion with feedback: the case of contact inhibition

Here we consider the case where a cell colony, spheroid or tumoroid, made of cells of a single type is subject to a feedback on its growth that depends on the population size. A common example is when proliferation rates depend on the local density on the population size. For instance, the release of promoting or inhibiting factors might increase (or decrease) the self-renewal probability. As another example, nutrient consumption or release of toxins , which increase with population size, might then in turn increase the probability of death.
We consider contact inhibition of growth as a reference biological situation[1]. In this situation, once confluence is reached, each cell will undergo several rounds of cell division while slowing down and will be reaching a regime where rare cell divisions are in equilibrium with rare cell deaths[2,3]. Cells can therefore grow with a self-renewal probability which decreases as the number of cells in the progeny increases, by increasing the probability of inactivation. In the following, we assume a simple linear functional form to define a BGW model capable of generating data with constrained dynamics, as sketched in fig. S4A. Mathematically, the functions governing the model parameters are defined as follows:

$$\begin{cases} \theta_0(Y) = \theta_0 \\ \theta_1(Y) = 1 - \theta_0 - \theta_2(1 - \frac{Y}{K}) \\ \theta_2(Y) = \theta_2(1 - \frac{Y}{K}) \end{cases} \tag{10}$$

where $K$ is a maximum clone size, constraining local density in the tissue, and $\vec{\theta}$ are analogous to the previous case. Inspired by experimental evidence on contact inhibition of growth in epithelial cell colonies[2], we set $K = 30$. This choice is motivated by the fact that cells can reduce their projected area of around a factor $K$ from when they fill the culture dish to when they arrest their growth. We performed numerical simulations with initial parameters $\vec{\theta} = (0.2, 0, 0.8)$, as shown in fig. S4B. Here Markovianity is assured by the fact that the probability of obtaining a population size $Y(n+1)$ starting from $Y(n)$ depends only on $Y(n)$ as in eqs. (10). Note that the precise choice of functional dependence is only needed to generate the synthetic data, but its knowledge is not required to perform the inference. Rather, it is the inference itself which reconstructs the functional form of the feedback.

All data generated from a given $Y$, regardless of the specific generation in which it occurs, can be grouped together by population size and used to infer the parameter values for that population size $\vec{\theta}(Y)$. The results of the inference performed with the EGS are shown in fig. S4C, where the mean and standard deviation of the marginals are displayed as a function of the clone size $Y$.

The comparison between the true and inferred parameter values clearly shows that our approach can be used to estimate the functional form of the feedback starting from data on the population size. Note that in all approaches presented here, inference is performed on data without tracking, assuming only the knowledge of the population size at all generations. Larger data-series can be handled by means of the MCEM approach which allows computationally more efficient inference. Results obtained by this method are shown in fig. S4D and highlight once more the capability of our approach to capture the true values with no prior knowledge.

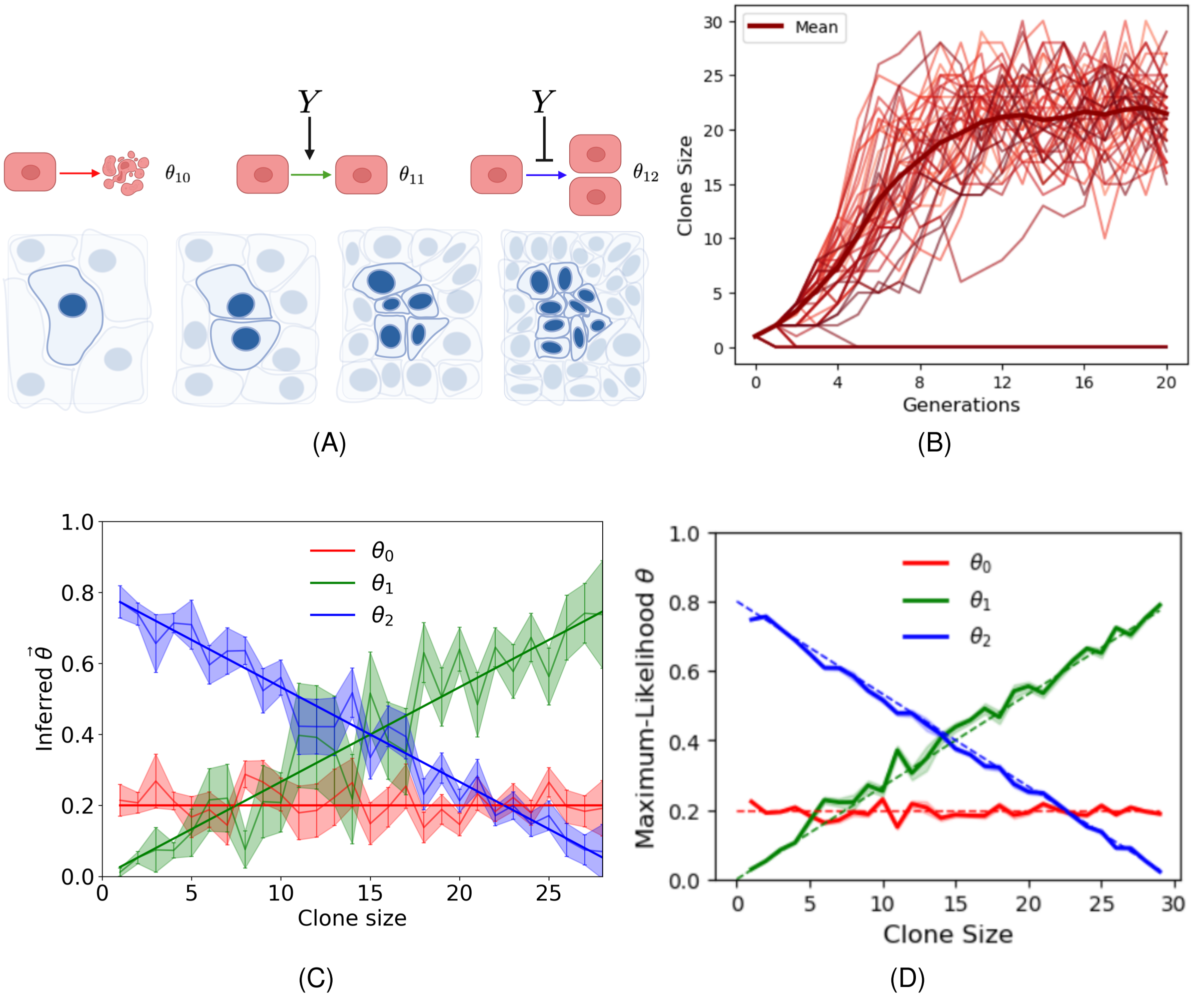

Figure S4: **Constrained expansion of cell clones A** (Top) Sketch of the constrained BGW model, where the total number of cells impacts on the probability of inactivity and proliferation. (Bottom) Illustration of the clonal growth in a confluent tissue, which represents the experimental counterpart of the data-series generated by numerically simulations. **B** Clone size as a function of the generation for the different realizations of the BGW process. Each clone starts from one cell, with parameters $\vec{\theta} = (0.2, 0, 0.8)$ and maximum clone size $K = 30$ cells. **C** Comparison between true and inferred parameter values obtained with the EGS: the true values are represented by dark solid lines, while the estimates are shown in lighter hues with colored regions indicating the corresponding standard deviation of the marginal distributions. **D** Comparison between true and inferred parameter values obtained with the MCEM method. True values are represented by dashed lines, while the estimates are shown in continuous lines.

### 4 MCEM convergence

A known limitation of MCEM methods is that they are local: when the likelihood function has multiple extrema, the algorithm may converge to a suboptimal solution corresponding to a local maximum rather than the global one. To mitigate this issue, one can run the algorithm multiple times with different initializations of the parameters, compute the likelihood at each MCEM step, and select the estimates corresponding to the highest likelihood found. An example is shown in Fig. S5, where the algorithm converges to two distinct sets of parameter values (corresponding to two different values of the likelihood). Computing the likelihood values (as shown in fig. S5) allows to rank the quality of estimates, even when the true parameter values are unknown.

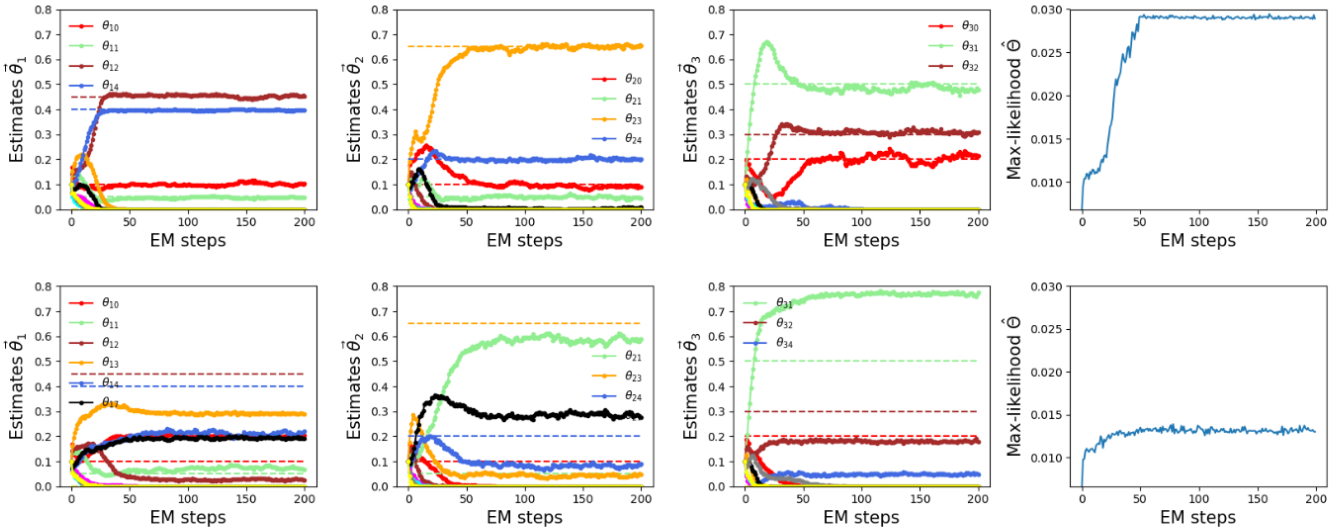

(A)

Figure S5: **Convergence of the MCEM** A three-type BGW process is used to perform inference. The number of parameters to infer is thirty (see main text). (Top) The inference converges to values close to the true ones. The right plot shows the corresponding value of the likelihood. (Bottom) A case where the same initial parameters, but different data samples, lead to a different set of inferred parameters which however correspond to a lower value of the likelihood.

### 5 Implementation of feedback on the total population for multi-lineage models

In this section we consider the effect of feedback on two different phenotypes. A possible biologically relevant scenario is the following. Suppose cells are able to degrade the medium or release certain inhibitory factors. Assume then that these processes inhibits the proliferation and the differentiation of stem cells while promoting quiescence and suppressing reversal to proliferative phenotype for differentiated cells, as shown in fig. S6A. We can model this situation with the following two-type BGW probabilities

$$\begin{cases} \theta_{10}(Y) = \theta_{10} \\ \theta_{11}(Y) = 0 \\ \theta_{12}(Y) = 1 - \theta_{10} - \theta_{13}(Y) \\ \theta_{13}(Y) = \theta_{13}\left(1 - \dfrac{Y}{K}\right) \end{cases} \qquad \begin{cases} \theta_{20}(Y) = \theta_{20} \\ \theta_{21}(Y) = 1 - \theta_{21} - \theta_{22}(Y) \\ \theta_{22}(Y) = \theta_{22}\left(1 - \dfrac{Y}{K}\right) \end{cases} \tag{11}$$

This model leads to a constrained growth in both subpopulations as shown in fig. S6B and promotes a change of abundance of cell of the two types, displaying stem cell dominated growth when $Y$ is small and viceversa when $Y$ is large. In order to perform inference with this model we generated $1000$ simulations starting with one stem cell. Results of the inference are shown in fig. S6D together with a comparison with the true parameters.

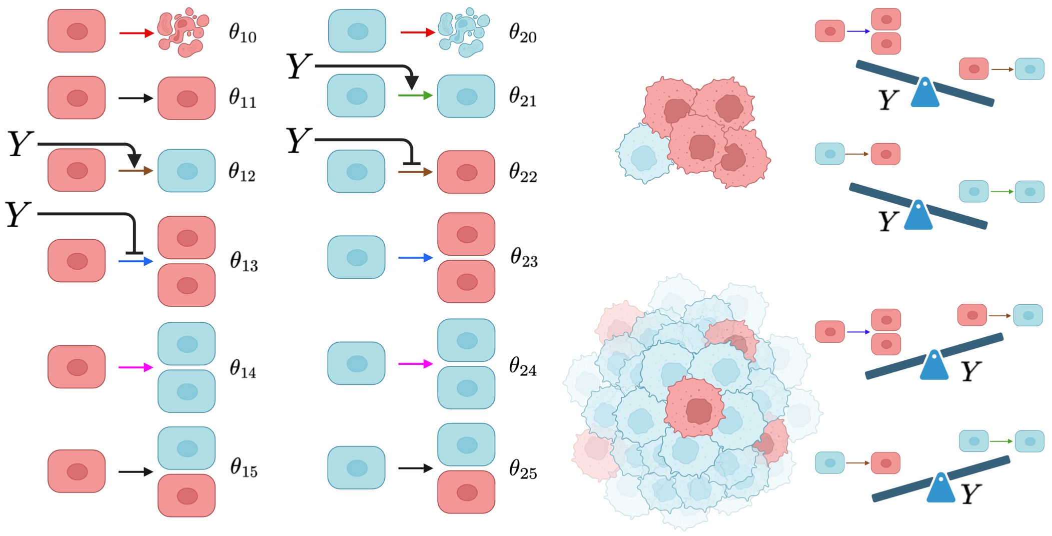

(A)

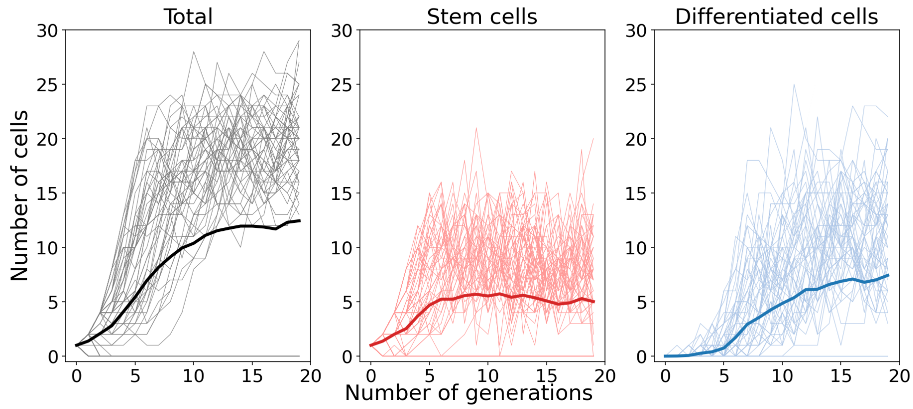

(B)

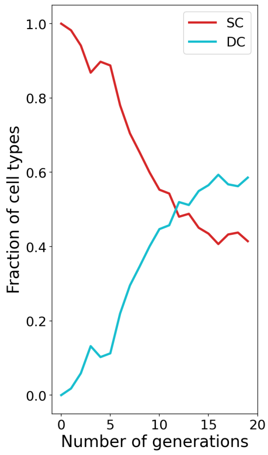

(C)

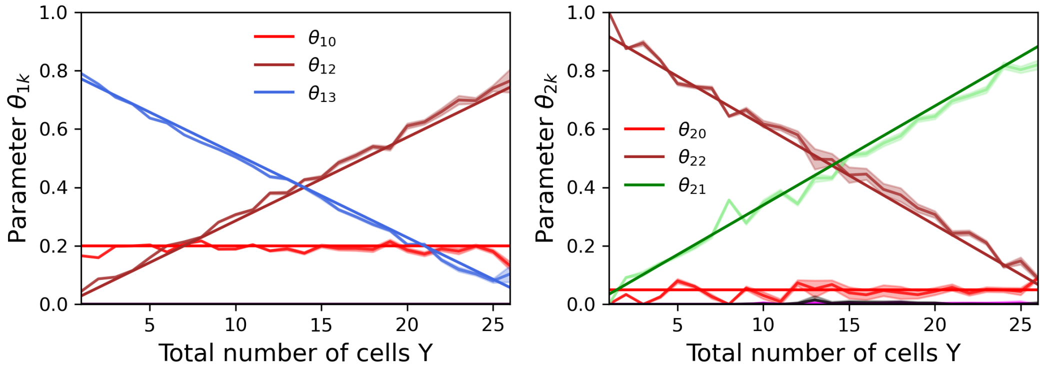

(D)

Figure S6: **Feedback on total population size A** Sketch of the model used with interactions modeled with a feedback on the total number of cells $Y$. The feedback is such that higher number of cells favor some transitions and inhibit others, as sketched. **B,C** Numerical simulations of the process shown in A, along with the fraction of stem vs differentiated cells which is changing over time. **D** Inference of parameters for stem cells (left) and differentiated cells (right) as a function of the total population size $Y$.

### 6 Inferring from non-clonal or asynchronous experimental data

Here we aim at testing our method against data where generations are not perfectly synchronous, for which the BGW branching process is only an approximation. Deviations from synchrony may take place because the initial condition is made of several non synchronized cells, or because division times are highly variable. In the first case, if the number of cells is very large, one might prefer to adopt a different model than BGW from the outset.

Clonal dynamics, i.e. the growth of small cell groups or single cells developing into a clone, is well suited to our approach as doubling times in cell populations are empirically found to be distributed according to Erlang distribution with relatively large shape parameters $k$[2,4–6]. Similar results have been shown for embryos[7,8], while these distributions are significantly more dispersed in bacterial populations[9]. For narrow doubling time distributions, the typical time for which the offspring remains significantly correlated is several generations or doubling times.

In order to show the performance of our method in this setting, we generated several numerical simulations with different doubling time statistics, i.e. generated with Erlang distribution with shape parameter $k = 10, 50, 100$, and rate parameters $\tilde{\lambda}k$ so that the mean doubling time mean is $\mu = \tilde{\lambda} = 1$, as shown in fig. S7. The relative dispersion for such distributions would be:

$$\frac{\sqrt{\sigma^2}}{\mu} = \frac{\sqrt{\frac{k}{\tilde{\lambda}^2 k^2}}}{\tilde{\lambda}k} = \frac{1}{\sqrt{k}} \tag{12}$$

where $k$ directly controls correlation between siblings (with $k = 1$ being the Poissonian case and $k \to \infty$ representing BGW). Numerical simulations of these birth-death processes were obtained by implementing Gillespie's method[10]. Cell death was simulated with a Poissonian process with rate $\delta$ (in the absence of more refined hypotheses), while cell proliferation would occur after $k$ Poissonian steps of rate $\lambda$, yielding an Erlang distribution for doubling time with shape parameter $k$ and rate parameter $k\tilde{\lambda}$.To demonstrate the effectiveness of our approach, we conducted tests using two different parameter sets, keeping the cell's duplication time equal to 1. In the first series, we assumed no cell death (fig. S7A), while in the second we enforced an approximate division probability per generation of $0.9$ and a death probability per generation of $0.1$ (fig. S7B). To carry out inference on the growth curves, we began by sampling the data under the imposed doubling time $\mu = 1$. We then aligned the trajectories to avoid ill-posed configurations, such as multiple consecutive divisions within a single generation time. To achieve this, we used numerically generated sequences of the form $[t, Y(t)]$ and applied a mean smoothing filter with a window size of $0.35\mu$. This allowed us to identify contiguous regions with non-positive derivatives. Among these, we decided to discard those shorter than $35\%$ of the doubling time.
We then searched for an appropriate time offset of the data such that sampling times $nT_{div}$ with ($n = 1, 2, 3$) would fall within the non positive derivative regions. Finally, we inferred transition probabilities using this sampled dataset.
For shape parameters and death rates similar to those encountered in experimental data, inference shows good results in both considered cases, as shown in fig. S7A,B. Our results indicate that deviations between true and inferred values lie within a range of 10% even considering asynchronous data closer to Poisson processes. Note that uncertainty is present also when data series are extracted from BGW processes as the amount of data is finite.
As expected, when lowering the shape parameters towards Poissonian dynamics inference is less and less reliable. One issue that is encountered is that different generations can overlap when cells become asynchronous, resulting in configurations that are incompatible with the model (for example observing more than $2^n$ cells at generation $n$). These cases can be handled in different ways, but would technically correspond to null likelihood and should therefore

be discarded. In general this issue is present whenever within one generation cells perform two consecutive transitions (for example two divisions or one division and one death). Cell death can easily be counted as happening at the next generation or, different definition of generation time can be chosen in order to solve this issue, for example half the division time. Another possibility is to introduce dedicated cell types which die with probability one. This new type, while fictitious and not directly measurable biologically, can introduce the kind of correlated behavior observed in experimental data, where generally the probability of cell death is very low but often happening in daughter cells.

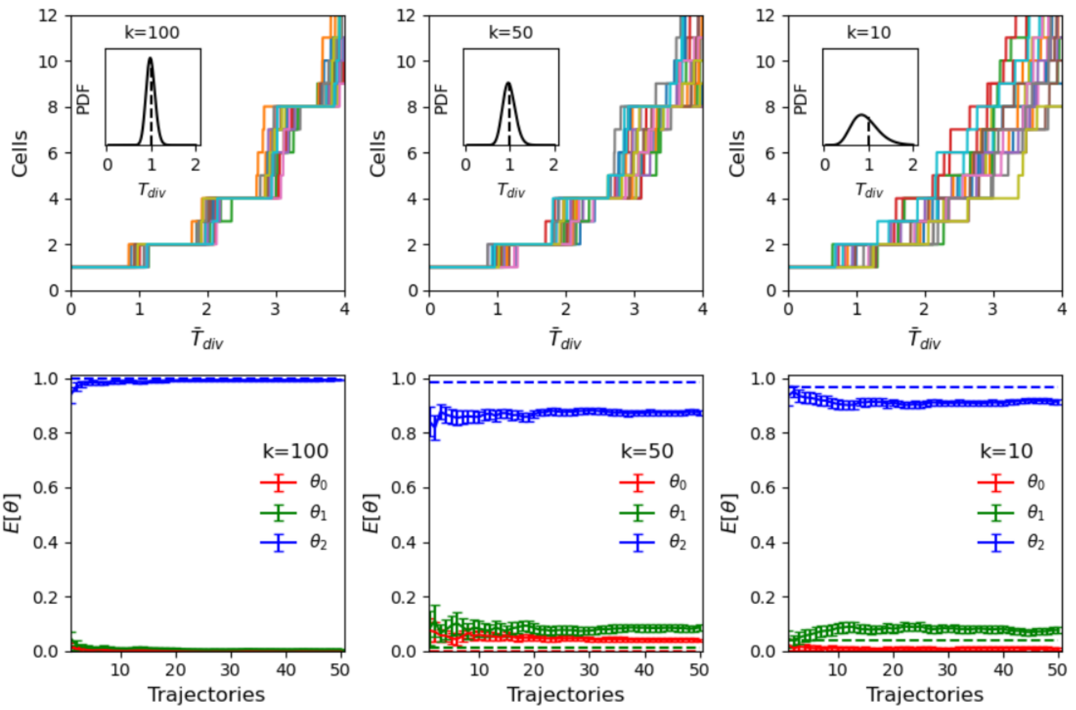

(A)

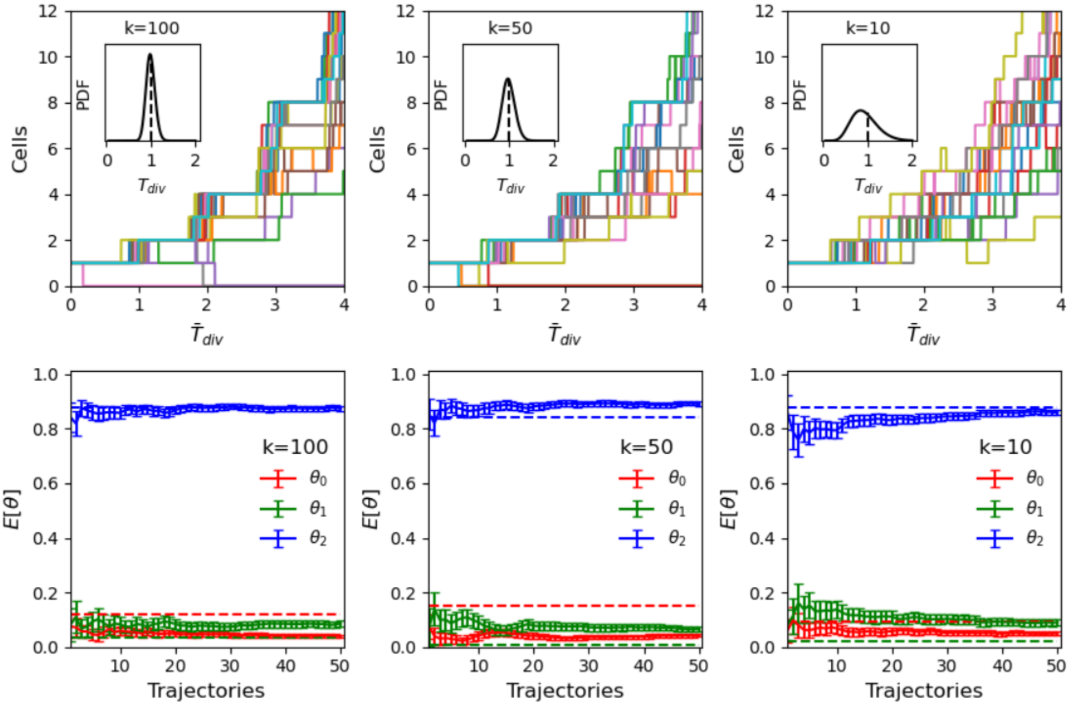

(B)

Figure S7: **Using BGW inference with asynchronous generation data. A** Numerical simulations of a birth process with cell doubling times distributed according to an Erlang distributions with shape parameters $k = 100, 50, 10$. Inference performed on $50$ trajectories $6$ generations long. **B** Numerical simulations of a birth-death process with cell doubling times distributed according to an Erlang distributions with shape parameters $k = 100, 50, 10$ and Poissonian deaths. Inference performed on $50$ trajectories $6$ generations long.

### 7 Monte Carlo Expectation Minimization with variable initial number of cells

The MCEM algorithm presented in the main text implicitly assumes that all observations at the $n-$th generation originate from the same initial number of cells. Here, we generalize the approach to accommodate observations derived from different initial cell counts.
We consider an initial number of cells denoted by $\mathbf{Y_i}$ and end up with a final number of cells $\mathbf{Y_f}$. Now, the likelihood explicitly contains the dependence from the initial population $\mathbf{Y_i}$:

$$\mathcal{L}(\mathbf{Y_f}|\mathbf{Y_i},\mathbf{\Theta}) = \sum_{\mathbf{Z}\in\{\mathbf{Z'}\}} p(\mathbf{Y_f}|\mathbf{Y_i},\mathbf{Z})p(\mathbf{Z}|\mathbf{Y_i},\mathbf{\Theta}) \tag{13}$$

When we perform the maximum-likelihood approach we need to sum also over the different initial conditions:

$$\hat{\mathbf{\Theta}} = \arg\max_{\Theta} \sum_{\vec{y}_f}\sum_{\vec{y}_i} \mu(\vec{y}_f,\vec{y}_i) \log \sum_{\mathbf{Z}\in\{\mathbf{Z'}\}} p(\mathbf{Y}_f|\mathbf{Y}_i,\mathbf{Z})p(\mathbf{Z}|\mathbf{Y_i},\mathbf{\Theta}) \tag{14}$$

where $\mu(\vec{y}_f,\vec{y}_i)$ is the joint empirical distribution that can be written as:

$$\mu(\vec{y}_f,\vec{y}_i) = \mu(\vec{y}_f|\vec{y}_i)\mu(\vec{y}_i) = \frac{1}{N_{\vec{y}_i}}\sum_{\vec{y}_f} I(\mathbf{Y}_f=\vec{y}_f|\mathbf{Y}_i=\vec{y}_i)\frac{1}{N}\sum_{\vec{y}_i} I(\mathbf{Y}_i=\vec{y}_i) \tag{15}$$

with $N_{\vec{y}_i}$ representing the number of observations starting with the initial population $\vec{y}_i$. Then we can write the surrogate function

$$\mathcal{B}(\mathbf{\Theta}) = \sum_{\vec{y}_f,\vec{y}_i,\mathbf{Z}\in\{\mathbf{Z'}\}} \mu(\vec{y}_f,\vec{y}_i)p(\mathbf{Z}|\vec{y}_f,\vec{y}_i,\mathbf{\Theta}) \log \frac{p(\mathbf{Y}_f|\mathbf{Y}_i,\mathbf{Z})p(\mathbf{Z}|\mathbf{Y_i},\mathbf{\Theta})}{p(\mathbf{Z}|\vec{y}_f,\vec{y}_i,\mathbf{\Theta})} \tag{16}$$

The maximum-likelihood estimate of each parameter becomes:

$$\hat{\theta}_{mt} = \frac{\sum_{\vec{y}_f}\sum_{\vec{y}_i}\mu(\vec{y}_i,\vec{y}_f)\sum_{\mathbf{Z}}p(\mathbf{Z}|\vec{y}_f,\vec{y}_i,\mathbf{\Theta})z_{mt}}{\sum_{t=0}^{T}\sum_{\vec{y}_f}\sum_{\vec{y}_i}\mu(\vec{y}_i,\vec{y}_f)\sum_{\mathbf{Z}}p(\mathbf{Z}|\vec{y}_f,\vec{y}_i,\mathbf{\Theta})z_{mt}} \tag{17}$$

Equation (23) in the main text is therefore replaced by:

$$\hat{Q}_{mt} = \frac{1}{L}\sum_{l=1}^{L} z_{ml}^{(l)}\tau_l(\mathbf{Y}_f^{(l)}=\vec{y}_f|\mathbf{Y}_i^{(l)}=\vec{y}_i)\mu(\mathbf{Y}_f^{(l)}=\vec{y}_f|\mathbf{Y}_i^{(l)}=\vec{y}_i)\mu(\mathbf{Y}_i^{(l)}=\vec{y}_i) \tag{18}$$

where the average is performed over different $L$ simulations each indicated by $l$, and $\tau_l(\mathbf{Y}_f^{(l)} = \vec{y}_f|\mathbf{Y}_i^{(l)}=\vec{y}_i)$ represents inter-arrival time between two identical observations that start from the same initial condition:

$$\tau_{l+1}(\vec{y}_f|\vec{y}_i) = \tau_l(\vec{y}_f|\vec{y}_i)\mathbf{I}(\mathbf{Y}_f^{(l)}\neq\vec{y}_f|\mathbf{Y}_i^{(l)}=\vec{y}_i)+1 \tag{19}$$

with initial condition $\tau_l(\vec{y}_f|\vec{y}_i) = 1$. With this estimator, each parameter $\theta_{mt}$ can be estimated as before through the formula

$$\hat{\theta}_{mt} = \frac{\hat{Q}_{mt}}{\sum_{t'}\hat{Q}_{mt'}} \tag{20}$$

This formulation generalizes the one presented in the main text and is needed for example in the case of multiple phenotypes with feedback, where there data series can start from different initial conditions, or in the case of experimental data.

### 8 Initialization strategies for MCEM inference

As previously pointed out, MCEM only guarantees convergence to local extreme and requires restart. The technique of moment matching allows to find a set of parameters that is able to give the same set of moments (and thus theoretically the same distribution/statistics) of a given random variable. This allows to select appropriate initial conditions for the parameters that avoid inconsistencies with the data. Note that for simple processes (for example the 1-type BGW) moment matching allows in principle to fully determine the parameters of the process starting from statistical observations such as mean, standard deviations and higher order moments. As detailed in the following, the situation for multi-type processes is far more complex. The approach below was followed for the inference presented in fig. 5H of the main text.

#### 8.1 Moments of a Multi-type Galton-Watson model

Let us start by considering a 3-types BGW model as described in the manuscript, characterized by the following PGFs:

$$\begin{aligned} f_1(s_1,s_2,s_3) =&\theta_{10}+\theta_{11}s_1+\theta_{12}s_2+\theta_{13}s_3+\theta_{14}s_1^2+\theta_{15}s_1s_2+\\ &+\theta_{16}s_1s_3+\theta_{17}s_2^2+\theta_{18}s_3^2+\theta_{19}s_2s_3 \end{aligned} \tag{21}$$

$$\begin{aligned} f_2(s_1,s_2,s_3) =&\theta_{20}+\theta_{21}s_2+\theta_{22}s_1+\theta_{23}s_3+\theta_{24}s_2^2+\theta_{25}s_2s_1+\\ &+\theta_{26}s_2s_3+\theta_{27}s_1^2+\theta_{28}s_3^2+\theta_{29}s_1s_3 \end{aligned} \tag{22}$$

$$\begin{aligned} f_3(s_1,s_2,s_3) =&\theta_{30}+\theta_{31}s_3+\theta_{32}s_1+\theta_{33}s_2+\theta_{34}s_3^2+\theta_{35}s_3s_1+\\ &+\theta_{36}s_3s_1+\theta_{37}s_1^2+\theta_{38}s_2^2+\theta_{39}s_1s_2 \end{aligned} \tag{23}$$

If we consider each type to start with a generic initial population $Y(0) = (Y_1, Y_2, Y_3)$, then the PGFs for the sub-populations can be written as

$$f_{Y_1}(s_1,s_2,s_3) = f_1(s_1,s_2,s_3)^{Y_1} \tag{24}$$
$$f_{Y_2}(s_1,s_2,s_3) = f_2(s_1,s_2,s_3)^{Y_2} \tag{25}$$
$$f_{Y_3}(s_1,s_2,s_3) = f_3(s_1,s_2,s_3)^{Y_3} \tag{26}$$

As detailed in the text, within BGW the random variable that describes the population dynamics is the random variable that contains the offspring of each lineage $m$ at the first generation, indicated with $\vec{X}_m(1) = \vec{X}_m = (X_{m,1}, \ldots, X_{m,M})$. The components $X_{m,i}$ represent the number of descendants of type $i$ generated by type $m$ in one generation, and where each index runs over $\{1, \ldots, 3\}$. The relationship between the number of cells of the $i-$th type and the offspring random variable $\{X_i\}$ is given by the equation:

$$Y_i(1) = \sum_{m=1}^{3} \sum_{k=1}^{Y_m(0)} X_{mi}^{(k)} \tag{27}$$

where the first sum is over the cell types, the second one is over the cells of a given type.
If one is interested in the moments of the process, it is convenient to work with the moment generating functions (rather than the PGFs), easily found from the PGFs through the substitution

$s_i \to e^{s_i}$:

$$\begin{aligned} M_1(\vec{s}) =& \theta_{10} + \theta_{11}e^{s_1} + \theta_{12}e^{s_2} + \theta_{13}e^{s_3} + \theta_{14}e^{2s_1} + \theta_{15}e^{s_1+s_2} + \\ & + \theta_{16}e^{s_1+s_3} + \theta_{17}e^{2s_2} + \theta_{18}e^{2s_3} + \theta_{19}e^{s_2+s_3} \end{aligned} \tag{28}$$

$$\begin{aligned} M_2(\vec{s}) =& \theta_{20} + \theta_{21}e^{s_2} + \theta_{22}e^{s_1} + \theta_{23}e^{s_3} + \theta_{24}e^{2s_2} + \theta_{25}e^{s_2+s_1} + \\ & + \theta_{26}e^{s_2+s_3} + \theta_{27}e^{2s_1} + \theta_{28}e^{2s_3} + \theta_{29}e^{s_1+s_3} \end{aligned} \tag{29}$$

$$\begin{aligned} M_3(\vec{s}) =& \theta_{30} + \theta_{31}e^{s_3} + \theta_{32}e^{s_1} + \theta_{33}e^{s_2} + \theta_{34}e^{2s_3} + \theta_{35}e^{s_3+s_1} + \\ & + \theta_{36}e^{s_3+s_1} + \theta_{37}e^{2s_1} + \theta_{38}e^{2s_2} + \theta_{39}e^{s_1+s_2} \end{aligned} \tag{30}$$

The **first moment** can be found through the relation

$$E[Y_i] = \sum_m \sum_k^{Y_m(0)} E[X_{mi}^{(k)}] = \sum_m Y_m(0) E[X_{mi}] = \sum_m Y_m(0) \frac{\partial M_m}{\partial s_i}|_{\vec{s}=\vec{0}} \tag{31}$$

We are also interested in the **second moment**:

$$E[Y_i(1)^2] = E[Y_i(1)Y_i(1)] = E\Bigg[ \Big( \sum_m^3 \sum_k^{Y_m(0)} X_{mi}^{(k)} \Big) \Big( \sum_{m'}^3 \sum_{k'}^{Y_{m'}(0)} X_{m'i}^{(k')} \Big) \Bigg] = \tag{32}$$

$$= E\Bigg[ \Big( \sum_m^3 \sum_{m'}^3 \sum_k^{Y_m(0)} \sum_{k'}^{Y_{m'}(0)} X_{mi}^{(k)} X_{m'i}^{(k')} \Big) \Bigg] = \tag{33}$$

$$= \sum_m^3 \sum_{m'}^3 \sum_k^{Y_m(0)} \sum_{k'}^{Y_{m'}(0)} E\Big[ X_{mi}^{(k)} X_{m'i}^{(k')} \Big] \tag{34}$$

There are 3 contributions in the above equation:

1. $m = m'$, $k = k'$: same type, same cell

$$E[X_{mi}^{(k)} X_{mi}^{(k)}] = E[X_{mi}^2] = \frac{\partial^2 M_m}{\partial s_i^2}|_{\vec{s}=\vec{0}} \tag{35}$$

2. $m = m'$, $k \neq k'$: same type, different cells

$$E[X_{mi}^{(k)} X_{mi}^{(k')}] = E[X_{mi}^{(k)}] E[X_{mi}^{(k')}] = \left( \frac{\partial M_m}{\partial s_i}|_{\vec{s}=\vec{0}} \right)^2 \tag{36}$$

3. $m \neq m'$, $k \neq k'$: different types, different cells

$$E[X_{mi}^{(k)} X_{m'i}^{(k')}] = E[X_{mi}^{(k)}] E[X_{m'i}^{(k')}] = \frac{\partial M_m}{\partial s_i}|_{\vec{s}=\vec{0}} \frac{\partial M_{m'}}{\partial s_i}|_{\vec{s}=\vec{0}} \tag{37}$$

Gathering all these terms together, we obtain:

$$\begin{aligned} E[Y_i(1)^2] =& \sum_m Y_m(0) \frac{\partial^2 M_m}{\partial s_i^2}|_{\vec{s}=\vec{0}} + \\ & + \sum_m Y_m(0) \Big( Y_m(0) - 1 \Big) \left( \frac{\partial M_m}{\partial s_i}|_{\vec{s}=\vec{0}} \right)^2 + \\ & + \sum_m \sum_{m'} Y_m(0) Y_{m'}(0) \frac{\partial M_m}{\partial s_i}|_{\vec{s}=\vec{0}} \frac{\partial M_{m'}}{\partial s_i}|_{\vec{s}=\vec{0}} \end{aligned}$$

In the same way we can obtain the **cross moments**:

$$\begin{aligned} E[Y_i(1)Y_j(1)] &= E\Big[\big(\sum_m^3 \sum_k^{Y_m(0)} X_{mi}^{(k)}\big)\big(\sum_{m'}^3 \sum_{k'}^{Y_{m'}(0)} X_{m'j}^{(k')}\big)\Big] = \\ &= E\Big[\sum_m^3 \sum_{m'}^3 \sum_k^{Y_m(0)} \sum_{k'}^{Y_{m'}(0)} X_{mi}^{(k)} X_{m'j}^{(k')}\Big] = \\ &= \sum_m^3 \sum_{m'}^3 \sum_k^{Y_m(0)} \sum_{k'}^{Y_{m'}(0)} E\Big[X_{mi}^{(k)} X_{m'j}^{(k')}\Big] \end{aligned}$$

There are again three contributions in the above equation:

1. $m = m'$, $k = k'$: same type, same cell

$$E[X_{mi}^{(k)} X_{mj}^{(k)}] = \frac{\partial^2 M_m}{\partial s_i \partial s_j}|_{\vec{s}=\vec{0}} \tag{38}$$

2. $m = m'$, $k \neq k'$: same type, different cells

$$E[X_{mi}^{(k)} X_{mj}^{(k')}] = E[X_{mi}^{(k)}]E[X_{mj}^{(k')}] = \frac{\partial M_m}{\partial s_i}|_{\vec{s}=\vec{0}} \frac{\partial M_m}{\partial s_j}|_{\vec{s}=\vec{0}} \tag{39}$$

3. $m \neq m'$, $k \neq k'$: different types, different cells

$$E[X_{mi}^{(k)} X_{m'j}^{(k')}] = E[X_{mi}^{(k)}]E[X_{m'j}^{(k')}] = \frac{\partial M_m}{\partial s_i}|_{\vec{s}=\vec{0}} \frac{\partial M_{m'}}{\partial s_j}|_{\vec{s}=\vec{0}} \tag{40}$$

Gathering all these terms together, we obtain:

$$\begin{aligned} E[Y_i(1)Y_j(1)] =& \sum_m Y_m(0) \frac{\partial^2 M_m}{\partial s_i \partial s_j}|_{\vec{s}=\vec{0}} + \\ &+ \sum_m Y_m(0)\big(Y_m(0)-1\big) \frac{\partial M_m}{\partial s_i}|_{\vec{s}=\vec{0}} \frac{\partial M_m}{\partial s_j}|_{\vec{s}=\vec{0}} + \\ &+ \sum_m \sum_{m'} Y_m(0) Y_{m'}(0) \frac{\partial M_m}{\partial s_i}|_{\vec{s}=\vec{0}} \frac{\partial M_{m'}}{\partial s_j}|_{\vec{s}=\vec{0}} \end{aligned}$$

At this point, we only need to evaluate the terms containing the derivatives of the mgfs:

$$E[X_{mi} X_{mj}] = \frac{\partial M_{X_m}}{\partial s_i \partial s_j}|_{\vec{s}=\vec{0}} \tag{41}$$

When $i \neq j$ these are:

$$\begin{aligned} &E[X_{11}X_{12}] = \theta_{15} \quad E[X_{11}X_{13}] = \theta_{16} \quad E[X_{12}X_{13}] = \theta_{19} \\ &E[X_{21}X_{22}] = \theta_{25} \quad E[X_{21}X_{23}] = \theta_{26} \quad E[X_{22}X_{23}] = \theta_{29} \\ &E[X_{33}X_{31}] = \theta_{35} \quad E[X_{33}X_{32}] = \theta_{36} \quad E[X_{31}X_{32}] = \theta_{39} \end{aligned}$$

whereas when $i = j$ these values are:

$$
\begin{array}{lll}
E[X_{11}^2] = 2\theta_{14} + E[X_{11}] & E[X_{12}^2] = 2\theta_{17} + E[X_{12}] & E[X_{13}^2] = 2\theta_{18} + E[X_{13}] \\
E[X_{21}^2] = 2\theta_{27} + E[X_{21}] & E[X_{22}^2] = 2\theta_{24} + E[X_{22}] & E[X_{23}^2] = 2\theta_{28} + E[X_{23}] \\
E[X_{31}^2] = 2\theta_{37} + E[X_{31}] & E[X_{32}^2] = 2\theta_{38} + E[X_{32}] & E[X_{33}^2] = 2\theta_{34} + E[X_{33}]
\end{array}
$$

#### 8.2 Moment Matching

Now we want to perform moment matching, i.e. determining the parameters in such a way that the theoretically obtained moments (as a function of the parameters) correspond to those obtained empirically from data. The obtained solution will act as a starting point to infer the underlying parameters with MCEM. Note that, for simplicity, we wrote this paragraph thinking to the data shown in fig. 5 of the manuscript. Therefore, we suppose to have cell counts $Y$ at two consecutive generations $0$ and $1$, so that, in what follows, $Y(1)$ can be thought as the number of cells at one generation, knowing the number of cells at the previous generation.
Let us denote with $\langle\ \ \rangle$ the empirical average. We therefore have access to all of the following quantities:

$$\{\langle Y_m(1)\rangle, \langle Y_m^2(1)\rangle, \langle Y_m(1)Y_{m'}(1)\rangle, \dots\}$$

We then have a (non-linear) system of equations with the parameters as unknown:

$$
\begin{cases}
E[Y_m(1)] = \langle Y_m(1)\rangle \\
E[Y_m^2(1)] = \langle Y_m^2(1)\rangle \\
E[Y_m(1)Y_{m'}(1)] = \langle Y_m(1)Y_{m'}(1)\rangle \\
\text{Var}(Y_m(1)) = \langle Y_m^2(1)\rangle - \langle Y_m(1)\rangle^2 \\
E[Y_m^2(1)Y_{m'}^2(1)] = \langle Y_m(1)Y_{m'}(1)\rangle \\
\text{Cov}(Y_m(1)Y_{m'}(1)) = \langle Y_m(1)Y_{m'}(1)\rangle - \langle Y_m(1)\rangle\langle Y_{m'}(1)\rangle \\
\vdots
\end{cases}
\tag{42}
$$

These are 18 relations in total, to which a normalization condition for each phenotype can be added: $\sum_t \theta_{mt} = 1$. Therefore, eq. (42) represents a non-linear undetermined system of equations (21 conditions for 30 unknowns). In order to increase the number of equations, we considered datasets as being organized into different unique initial conditions:

$$\{(Y_{1,r}(0), Y_{2,r}(0), Y_{3,r}(0))\}$$

where each $r$ is an index for a different initial configuration. At this point, we can write 18 equations for each index $r$, and find (numerically) a solution eq. (42).
In order to find a set of parameters satisfying eq. (42), we resort to the numerical minimization

of the sum of the squares of the terms:

$$
\begin{aligned}
\min_{\Theta} \Bigg\{ \sum_r \Bigg[ & \sum_m (\sum_t \theta_{mt} - 1)^2 + \\
& + \sum_m \Big( E[Y_{m,r}(1)] - \langle Y_{m,r}(1) \rangle \Big)^2 + \\
& + \sum_m \Big( E[Y^2_{m,r}(1)] - \langle Y^2_{m,r}(1) \rangle \Big)^2 + \\
& + \sum_m \sum_{m' \neq m} \Big( E[Y_{m,r}(1) Y_{m',r}(1)] - \langle Y_{m,r}(1) Y_{m',r}(1) \rangle \Big)^2 + \\
& + \sum_m \Big( \text{Var}(Y_{m,r}(1)) - \langle Y^2_{m,r}(1) \rangle + \langle Y_{m,r}(1) \rangle^2 \Big)^2 + \\
& + \sum_m \sum_{m' \neq m} \Big( \text{Cov}(Y_{m,r}(1) Y_{m',r}(1)) - \langle Y_{m,r}(1) Y_{m',r}(1) \rangle + \langle Y_{m,r}(1) \rangle \langle Y_{m',r}(1) \rangle \Big)^2 \Bigg] \Bigg\}
\end{aligned}
\tag{43}
$$

where the minimization is performed over the entire set of parameters. Note however that once a minimum is found, it is still necessary to verify that it is a solution to the system.
In practice, we considered three different initial conditions:

$$
\begin{aligned}
Y_{1,1}(0) = 5, Y_{2,1}(0) = 32, Y_{3,1}(0) = 3 \\
Y_{1,2}(0) = 6, Y_{2,2}(0) = 30, Y_{3,2}(0) = 4 \\
Y_{1,3}(0) = 3, Y_{2,3}(0) = 27, Y_{3,3}(0) = 2
\end{aligned}
$$

Each of these initial conditions was used to generate via numerical simulations of BGW process a total of 1500 realizations with the given parameters. At this point, $1500 \times 3$ data series spanning one generation (one for each initial condition) were used to write $18 \times 3$ equations to minimize eq. (43). For each of these we obtained a value for the 30 parameters, as shown in fig. S8.

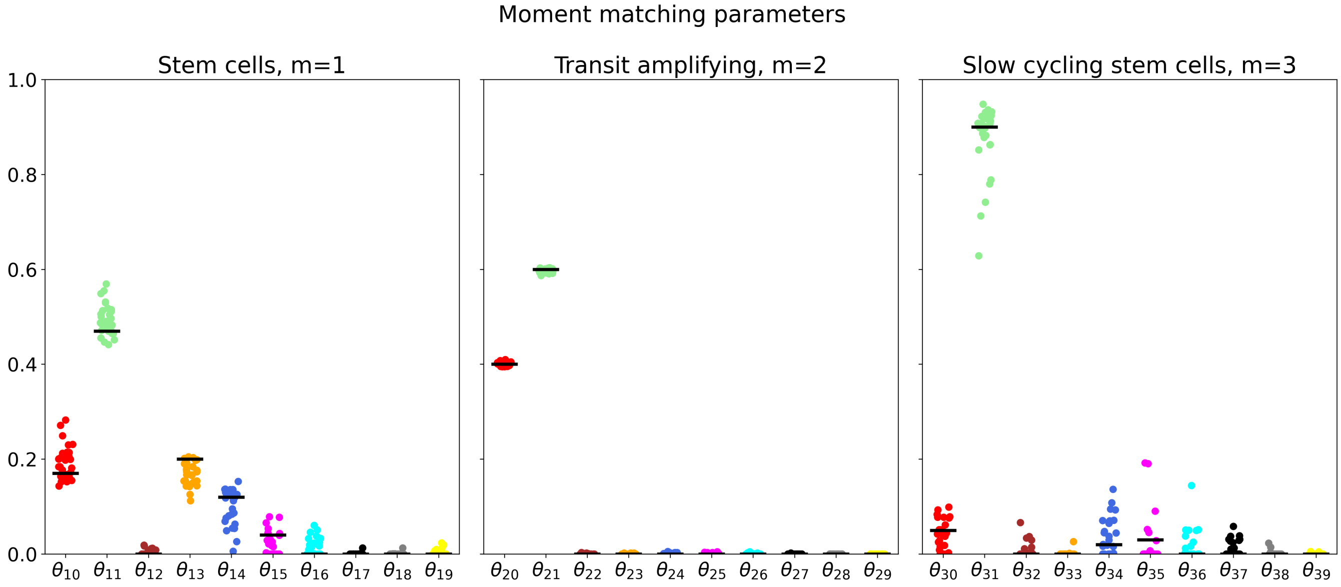

Figure S8: **Parameter estimation with moment matching technique.** Values of parameters obtained (colored points) for each parameter by means of moment matching explained in the text. The corresponding true value is plotted as a black dash. Each solution is found by performing sub-sampling on data generated with numerical simulations with different initial conditions.

### 9 Supplementary figures

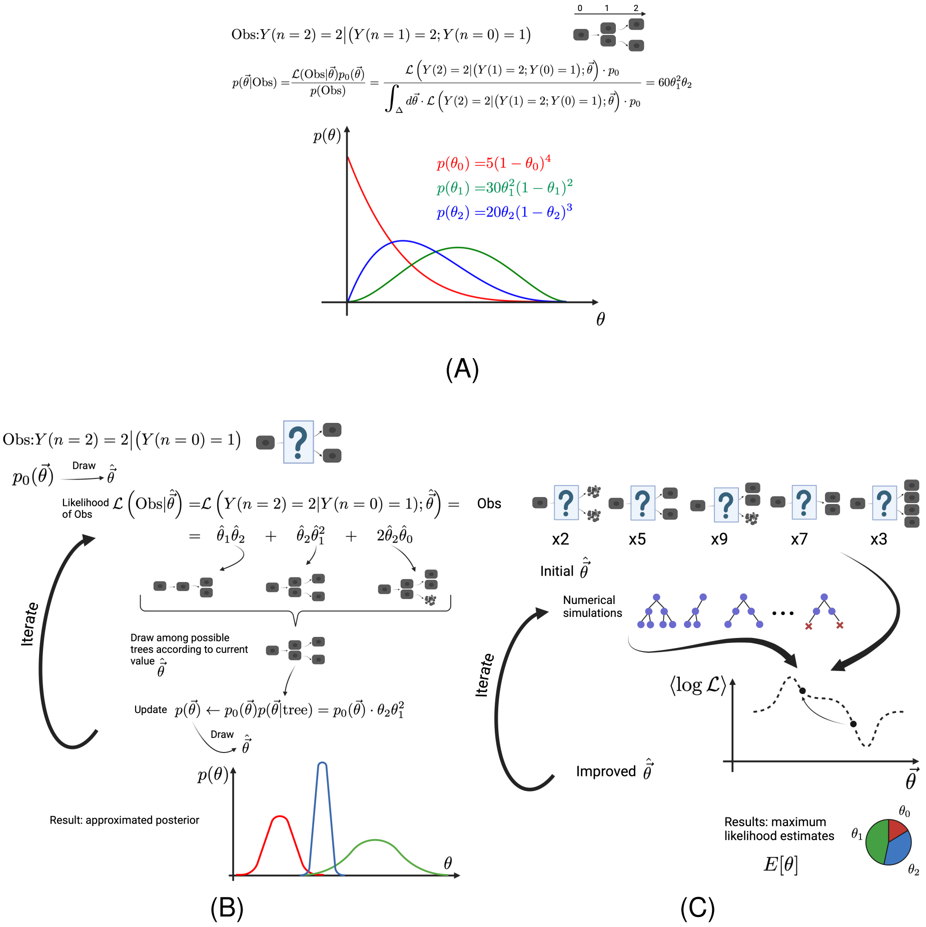

Figure S9: **Bienaymé Galton Watson branching processes and inference for cell populations A** Sketch of the methodology to infer transition probabilities with the analytical method. One full observation is assumed, along with associated transitions (one division and two inactivities). The corresponding posterior is explicitly worked out by means of Bayes' formula with a constant prior $p_0(\vec{\theta}) = p_0$. The normalization factor at the denominator is integrated over the parameters simplex (in this particular case a triangle). The posterior is then marginalized to obtain the analytical probability distribution functions for each parameter. Different observation are all factorized into the likelihood and processed in the same way. **B** Sketch of the methodology to infer transition probabilities with the EGS method. With a given observation (which here does not include the number of cells at the first generation) one draws a set of parameters distributed according to the prior. This set of parameters is used to evaluate the probability of each possible tree compatible with the given observation as shown in the sketch. Once one of the compatible trees is drawn the distribution of parameters is updated accordingly, and is then used to draw a new set of values. The procedure is iterated to compute the full marginal PDF of each parameter. **C** Sketch of the methodology to infer transition probabilities with the MCEM method. A set of observations with unknown intermediate states is assumed, along with a multiplicity. A set of parameters is drawn and numerical simulations of the underlying process are performed. These simulations are weighted with the empirical distribution of the observations and contribute to the estimation of the likelihood. The local extrema of the likelihood are searched by means of a surrogate function. This procedure, once converged, outputs the maximum likelihood estimation of each parameter.

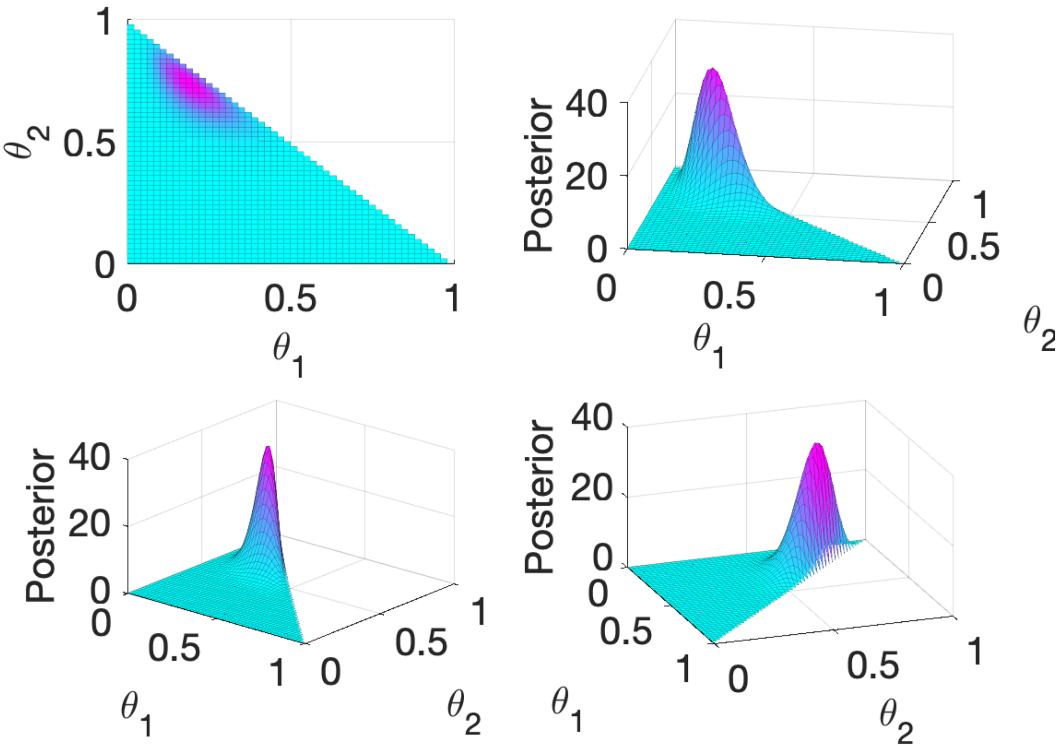

Figure S10: **Multivariate posterior.** The plots show different view of the multivariate posterior distribution obtained for the function $p(\mathbf{\Theta}|\mathbf{Y}) = \theta_0^2\theta_1^4\theta_2^{16}/\mathcal{N}$, where $\mathcal{N} = \Gamma(3)\Gamma(5)\Gamma(17)$.

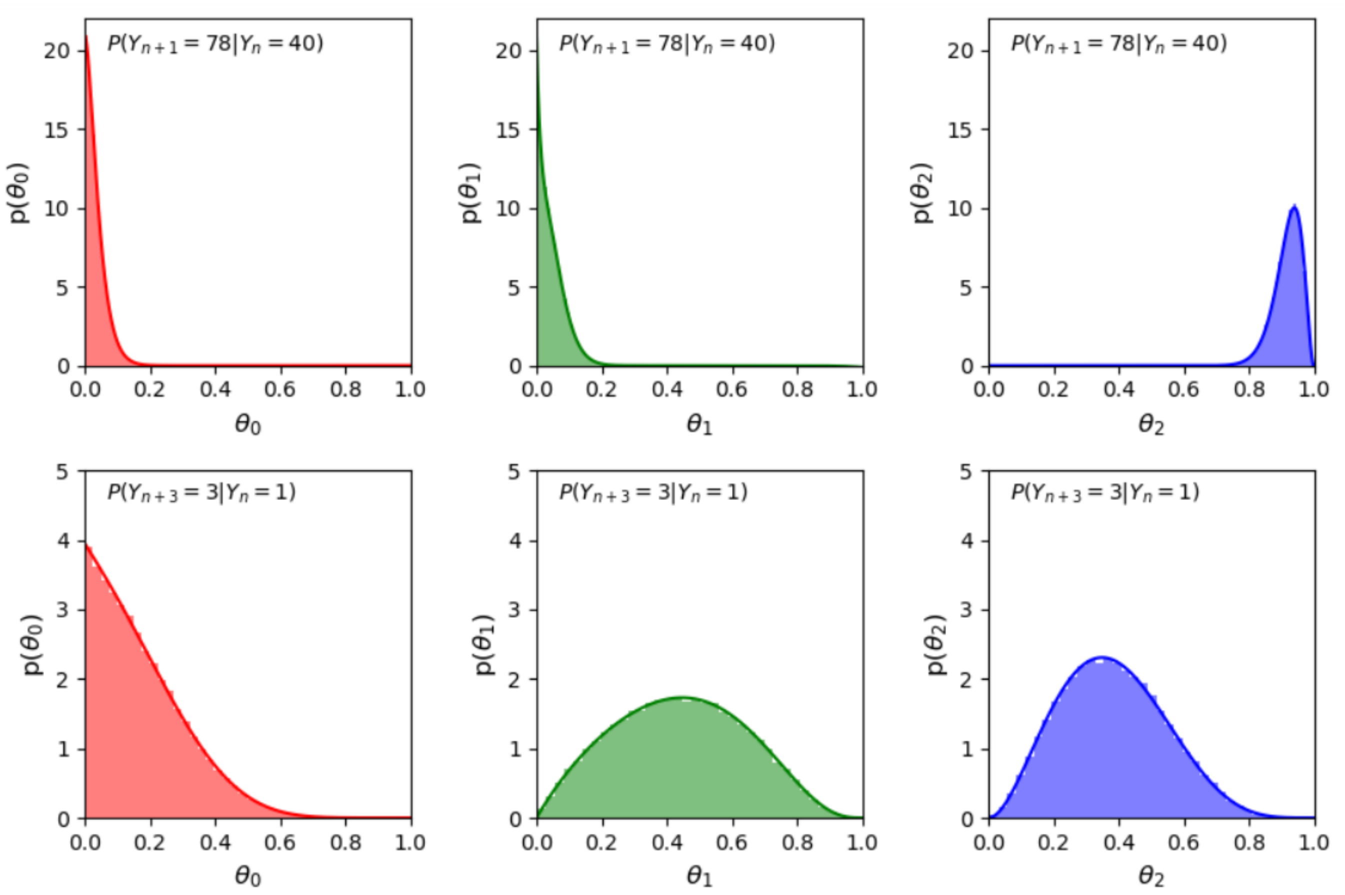

Figure S11: **Comparison between analytical and EGS marginals.** (Top) Marginals distributions for a BGW with one phenotype where $78$ cells are observed after one generation starting from $40$ cells. The marginal distributions obtained with EGS (in the lighter colors) are compared with the analytical (plotted in solid color) (Bottom) A case where $3$ cells are observed after 3 generations starting with a single cell is considered. A Kolmogorov-Smirnov two-tailed test for EGS marginals computed with $10^5$ points tested against discretized (100 points) exact marginal distributions yielded acceptance of the null hypothesis (the two come from equal distributions) at 5% significance level ($p-$value smaller than $10^{-6}$ for all marginals).

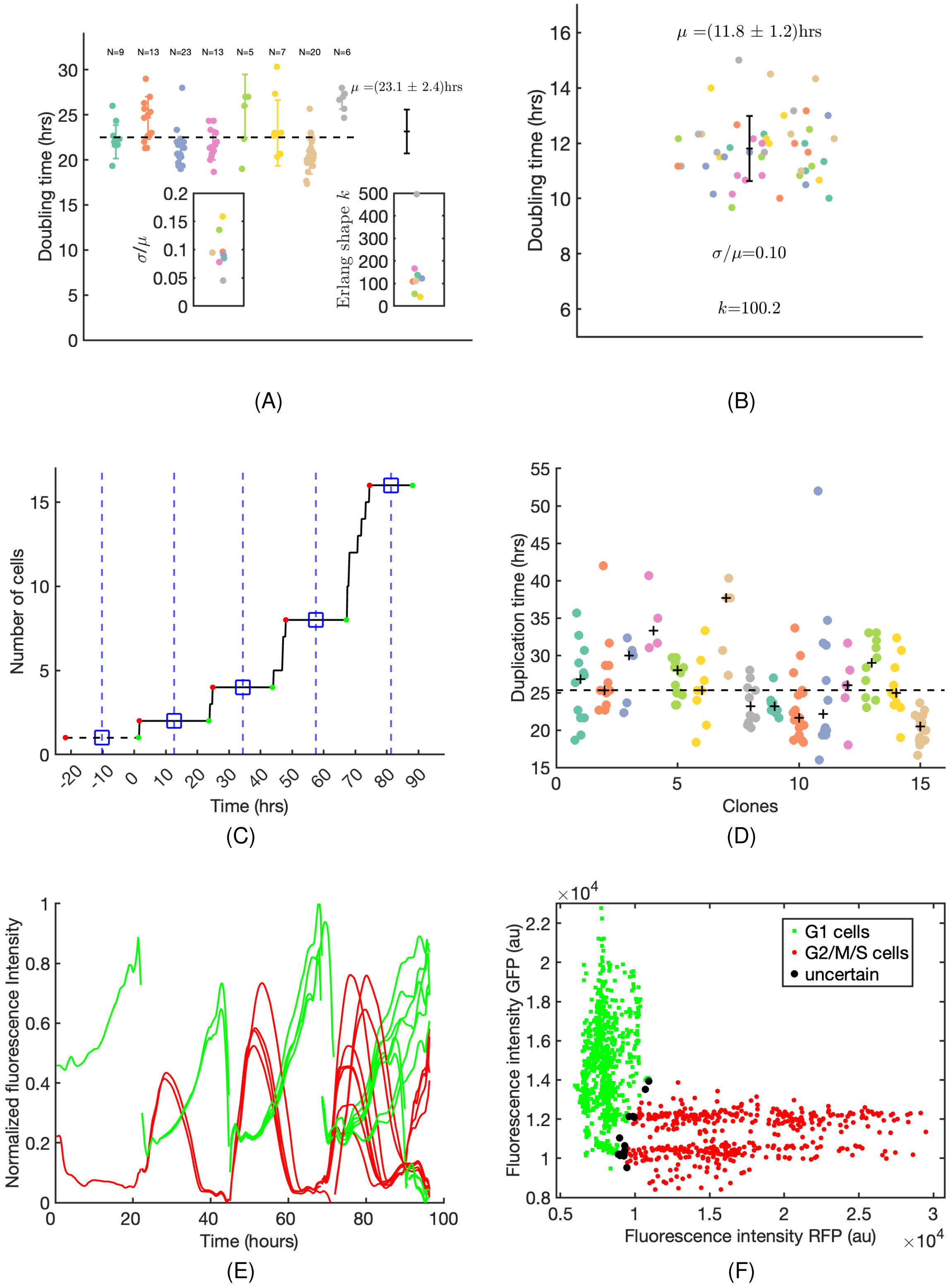

Figure S12: Caption is on the next page.

Figure S12: **A** Doubling times of offspring of single MG63 cells, relative to fig. 6A-F. Each cloud plot represents a clone and its offspring. The insets show the relative dispersion in each offspring and the corresponding shape parameter $k$ for an Erlang distribution. **B** Distribution of doubling times for MDCK cells relative to fig.6G-I in the initial free growth phase, along with relative dispersion and corresponding shape parameter for an equivalent Erlang distribution. **C** Distribution of duplication times for each clone relative to fig. 7. **D** Explanatory figure for experimental data processing. The plateaus in the original signal (black continuous line) are found along with start and end points (red and green dots respectively). Sampling points are found as those closest to plateau midpoints (blue empty squares), and spaced with the doubling time. Dashed blue lines correspond to the result of the minimization (see explanation on the METHODS section). **E** Fluorescent signal coming from single cells with a common single ancestor during a timelapse experiment. **F** Scatter plot of green vs red channel signal for a given clone in the timelapse.
